# Greenhouse gas flux and microbial communities in Central Valley irrigation canals

**DOI:** 10.1101/2025.07.03.662220

**Authors:** Mo Kaze, Roberto Sanchez, Mark Sistrom, Susannah Tringe

## Abstract

Climate change is of alarming and immediate concern and it is crucial to examine myriad facets contributing to the emission of greenhouse gases. Research into ecosystem microbial community structure and functional genetic diversity is essential for insight into global change and provides much-needed data for predictive models. Engineered transitional systems, such as aqueducts and networks of irrigation canals, have not been investigated for carbon flux and microbial community structure. Engineered interfaces may be acting as “hotspots” with impacts disproportionate to their geographic area. The Delta-Mendota aqueduct in California is 117 miles long and spans five counties; it is connected to 723 miles of irrigation canals within Merced County. This project combines field greenhouse gas measurements and culture-independent metagenomic analysis to identify the microbial communities of these two systems. This work maps the biogeographical distribution of organisms within these systems and defines their functional potential. DNA was extracted from 97 samples of sediment along 500 miles of irrigation canals in Merced County and sequenced with high-throughput whole genome sequencing. Each site was measured for methane and carbon dioxide flux with a field gas analyzer. Gas flux data was processed and visualized using multiple R packages. Metagenomic data was processed using multiple bioinformatics tools to bin and annotate genomes, normalize and quantify functional genes, and map the biogeographic distribution and phylogeny of identified microorganisms. Eighty-nine of the sample sites within the irrigation canal system demonstrated elevated carbon dioxide levels when compared to ambient air measurements (97%) and 43 sites had methane flux (44%). Results indicate a wide diversity of microorganisms present at each sampling site. Metagenomic analysis identified the microorganismal community of the sampled waterway sediments and confirmed the presence of metabolic pathways for methanogenesis and methane oxidation. Methanogenic genes were most abundant in methane producing sites. This community structure analysis reveals the potential of microbial biogeochemical activity and the gas measurements signal that greenhouse gas emission is currently occurring in these waterways. These results strongly suggest the need for monitoring and further investigation into other engineered terrestrial-aquatic interfaces.

## Introduction

Climate change is an alarming and immediate concern and to address it we need to examine all factors contributing to the emission of greenhouse gases. These dData are needed as input for predictive models to determine outcomes of global change, and identify priorities for mitigation.

There is well characterized dynamic carbon flux at interfaces between aquatic and terrestrial systems that impacts on global climate change.^1,2^ Many engineered transitional systems, such as aqueducts and networks of irrigation canals, have yet to be investigated for carbon flux.

Human-made waterway sediments likely contain microorganismal communities that impact carbon gas flux; mapping the biogeographical distribution of organisms and defining their functional potential across the water distribution system will provide input data for predictive models of landscape features not currently reflected in climate models.^9,10^

The Delta-Mendota aqueduct is 117 miles long and spans five counties in California (USGS). There are 723 miles of canals within the 154,000 total acres of the Merced Irrigation District Watershed (CA Water Board, 2013). These terrestrial aquatic interfaces cover large regions and are not currently being routinely measured for carbon flux and the microbial communities of these waterways and their potential contribution of greenhouse gases to the atmosphere has yet to be investigated. These engineered interfaces may act as “hotspots” with impacts disproportionate to their geographic area and there is evidence that this is the case in other systems.^11^ Identifying such hotspots within these large-scale engineered terrestrial aquatic habitats, and characterizing their resident microbial communities, may determine whetherif these interfaces require monitoring and mitigation are warranted.^12,13^ This pilot study investigated whether irrigation canals harbor specialized and the microorganismal communities adapted to these manmade environments that contribute to greenhouse gas cycling.

This study involved collecting 100 samples of sediment from the Delta-Mendota Aqueduct and Merced County irrigation canal system of the Central Valley of California. (Figure 1) Sediment brought to the surface was measured for carbon dioxide and methane flux. This pilot project examined the hypothesis that soil microbial communities in the sediment of waterways and canals originating from the San Joaquin Delta of California may be contributing greenhouses gases to the atmosphere. We founded two hypotheses based on the likely presence of run off from orchards, dairies, and feed lots abutting the irrigation canals, 1) there will be greenhouse gas emission from some of the canals and 2) methane emission would be associated with the presence of methanogenic species in the sediment’s microbial community. To test these hypotheses, we sampled sediment and took greenhouse gas measurements using a portable gas analyzer from one hundred different irrigation canal locations in the Central Valley of California in Merced County over ten dates during August and September of 2019. The sediment collections were sequenced with a culture-independent metagenomics approach using high-throughput whole metagenome sequencing to identify the microbial community members and their functional capabilities. We found methane and carbon dioxide flux occurring at many of the sampled locations and from the metagenomic data, identified methanogenic species and functional genes participating in methanogenesis pathways.

**Figure 1.**
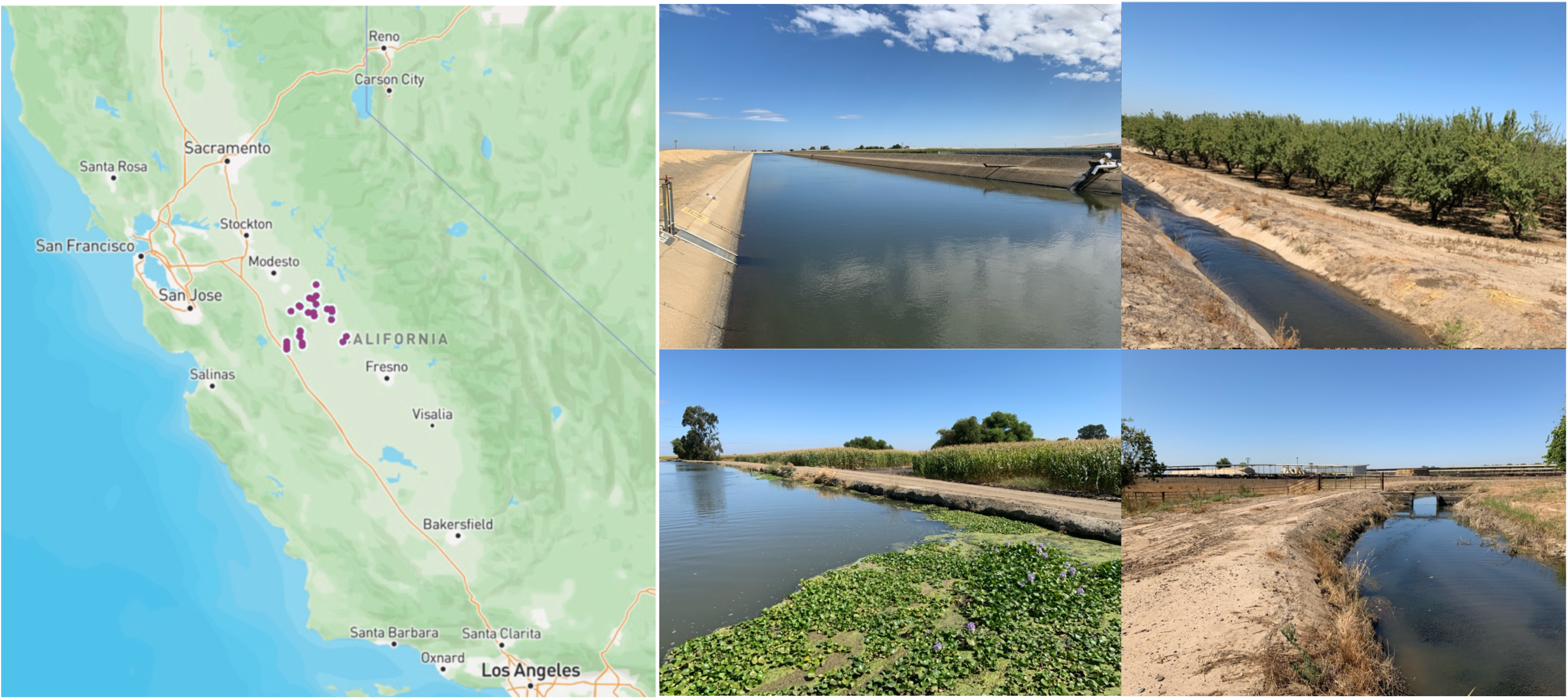
Location of sampling location and examples of engineered water movement systems in the Central Valley of California. A. Map of California where purple dots indicate sampling regions in Merced County. B.i. Delta-Mendota aqueduct. ii. Irrigation canal abutting almond orchard. iii. Wider irrigation canal with plant mat next to corn field. iv. Canal running next to large dairy (seen in background).

## Methods

### Sample locations and conditions

One hundred and ten sample locations were randomly selected using an ArcGIS map of the irrigation canal system of Merced County in the Central Valley of California. Ten locations were not sampled due to inaccessibility or safety concerns. Ninety-six irrigation canals and four sites along a large central aqueduct accessible from road shoulders were sampled.

Observations of soil pH were completed with test strips. Collected water temperature was measured with a standard analog thermometer. Flow rate was observed and qualitatively classified by researchers on a scale of stagnant (not moving), to moderate flow to rapid flow. Ambient field temperatures ranged from 99°F to 115°F over the sampling periods during July, August, and September. Sampling depth for collection was based on measurements taken from the distance markings of the ponar connector.

### Sample collection and processing

Samples were dredged from the floors of each irrigation and aqueduct sample site using a standard Ekman grab ponar. The ponar was aimed toward the center of each canal and the aqueduct. Sediment, measured out to 0.25L, was placed in a 0.5L glass mason jar and connected to the field gas analyzer system. Measurements were taken for 6 minutes total.

Samples were capped tightly and placed on ice. One gram of sediment was used for DNA extraction using DNeasy Powersoil Kit (Qiagen, Germany).

### Bioinformatics & Statistics

Files were converted from text documents to comma separated value documents and given column headers and each row provided the associated sampling site ID in R and cleaned for readability and transformed into matrices for analysis using fs, readr, and tidyverse packages.^14,15,16^ All raw and formatted output is available in the associated supplemental documentation. Initial bioinformatic processing and output was performed and provided by JGI (REF); taxonomic output from JGI was rounded to the nearest whole value and filtered for results with percent identity greater than or equal to 90% and 80% and greater for binned metagenomic data. Quantifications and visualizations were performed in R with package ggplot2 and image editing done in Preview.app (Apple Inc, Cupertino, CA).^17^ Sampling and sites where flux was measured were visualized with R packages mapview, sf, ggmap, and stamen.^18,19,20,21^ Color palettes were provided by R package viridis.^22^ Statistical analysis, Pearson and Spearman correlations were computed with base R and visualized using GGally and ggpubr’s function corr.test.^23,24^

Gas flux was calculated using the first 60 seconds of gas measurement in base R to identify the slope of the line (coef function) and by R packages gasfluxes and data.table.^25^ Initial gas measurements were converted from ppm to mg/m3 with equation mg/m3 = (X ppm)(molecular weight)/24.45. Data were processed and greenhouse gas flux was calculated using the non-linear HMR modeling method in gasfluxes R package.^26^Data normalization and calculation of alpha and beta diversity indices were performed using R package vegan.^27^ Visualization was also completed using gasfluxes output function: plot(). Analysis was double checked for consistency using R linear modeling package FluxCalR.^28^ We focused on four archaeal methanogenesis pathways using different methanogenic substrates (KEGG modules M00356 (methanol), M00357 (acetate), M00563 (methylamine/dimethylamine/trimethylamine), and M00567 (CO_2_ and H_2_).^29^

## Results

### Methane and Carbon Dioxide Flux from Irrigation Canal Sediment

Methane and carbon dioxide fluxes were measured from sediment dredged up from the bottom of one hundred sampled irrigation canals. Thirty-eight of the 100 sampling locations had methane production above background. Ninety-one sampling locations had carbon dioxide flux. (Figure 2) The majority of the methane flux occurred at locations in the Merced rural region with 11 locations (30%) and in nine locations in Dos Palos with (24%). Data were processed and greenhouse gas flux was calculated using the HMR method in R; methane fluxes ranged from 0.3 to 26.6 with a mean of 3.78, a median of 2.2, and a standard deviation of 5.3. (Figure 3) There were 22 sites with CO_2_ flux in the Dos Palos region and 20 sites with flux in Winton region. The range for carbon dioxide fluxe calculations was 0.54 to 408.2 with a mean of 34.5, a median of 14.1 and standard deviation of 62.1. (Figure 4) Samples from six locations in Hilmar, Livingston, and Winton, had no measured gas emission. (Table 1). Assessing correlation of methane flux to carbon dioxide flux, we found there was positive correlation between the methane and carbon dioxide flux (Pearson, 0.45, p ≤0.5). (Sup Figure 1A) Each site that produced methane also produced carbon dioxide, however, the converse was not true. (Sup Figure 1A)

**Table 1.**
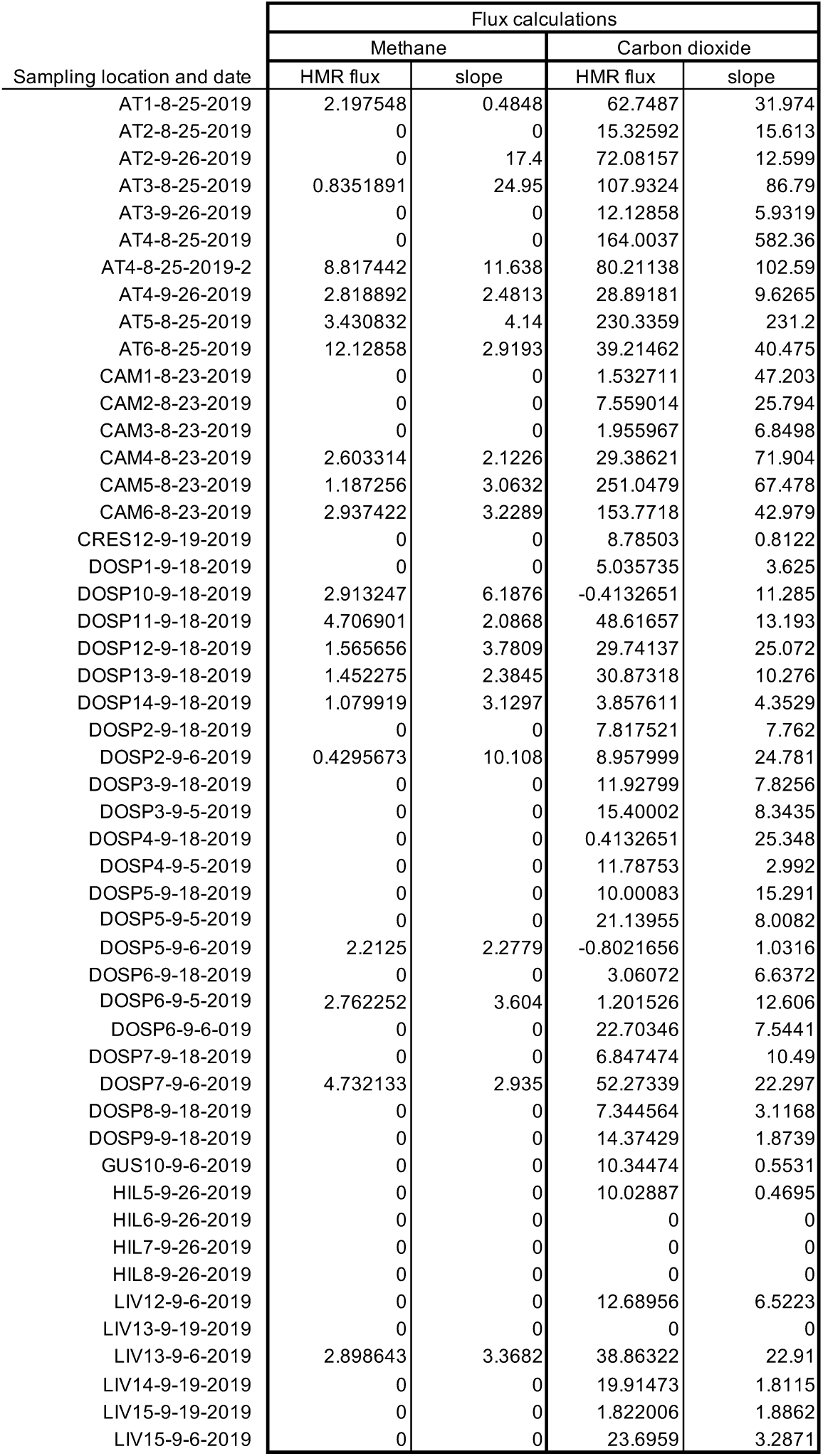
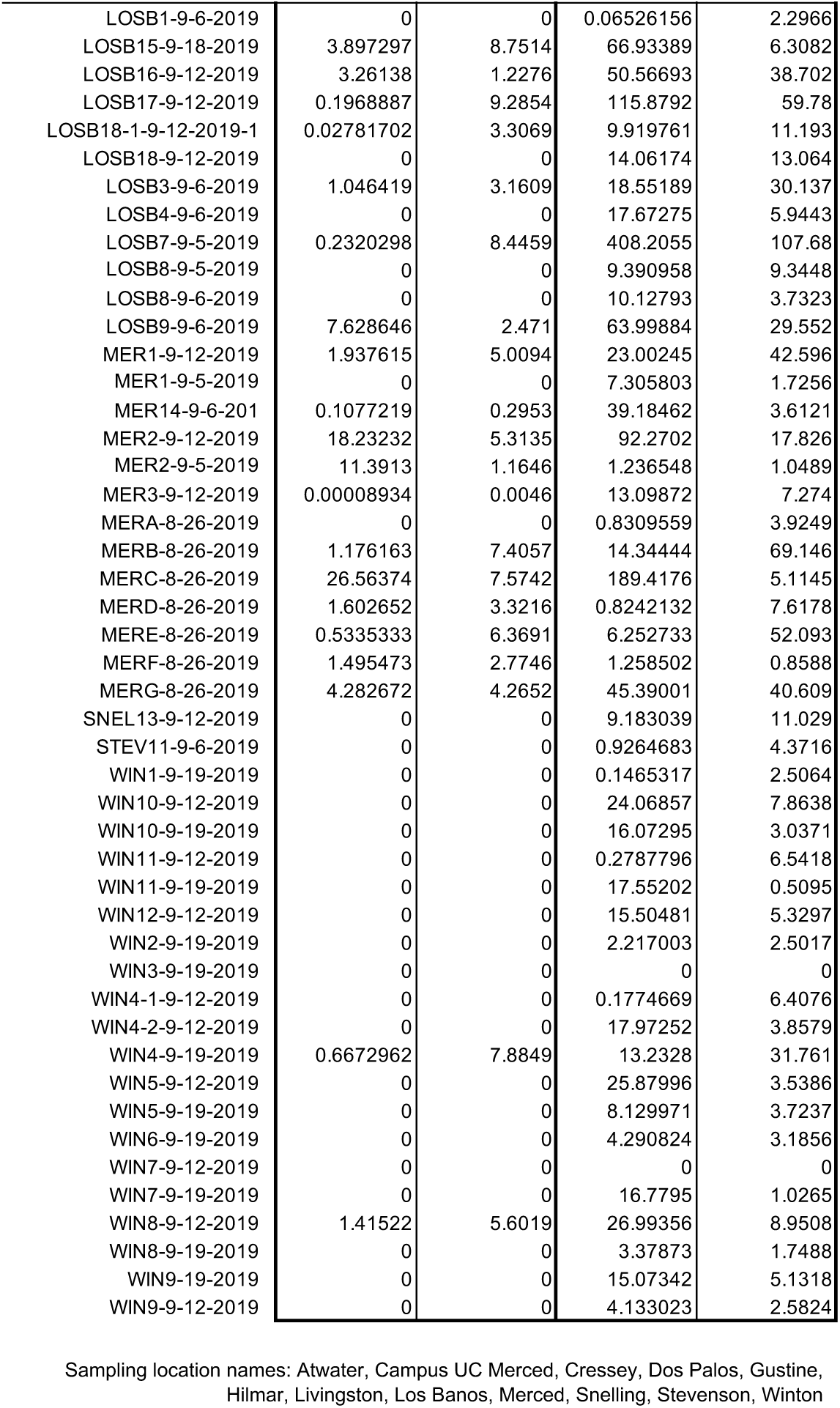
Calculated methane and carbon dioxide flux for ninety-seven sampling locations.

**Figure 2.**
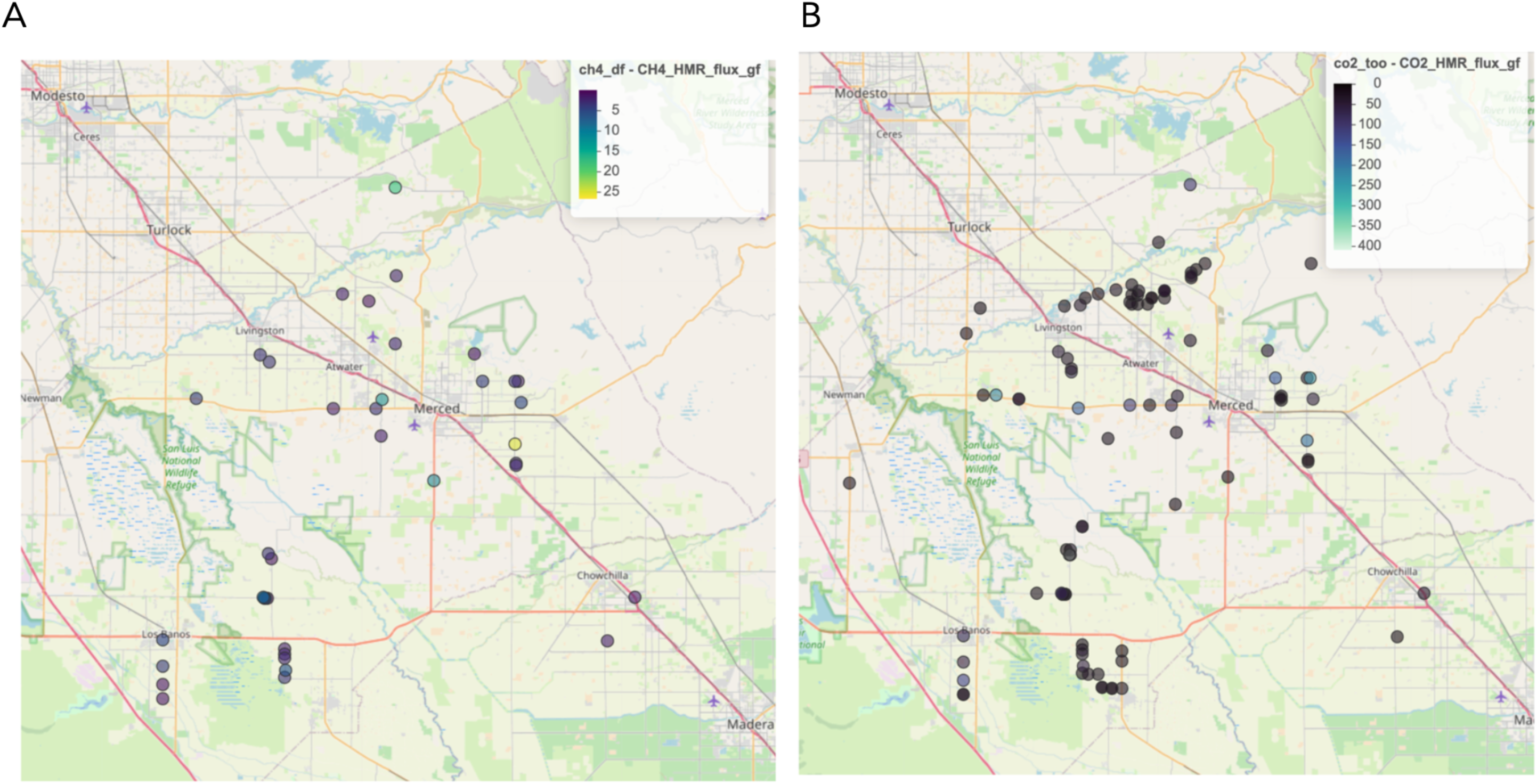
Greenhouse gas flux results. A. Methane flux occurred at 38 of the 100 sampled locations. B. Carbon dioxide flux occurred at 91 of the 100 sampled locations.

**Figure 3.**
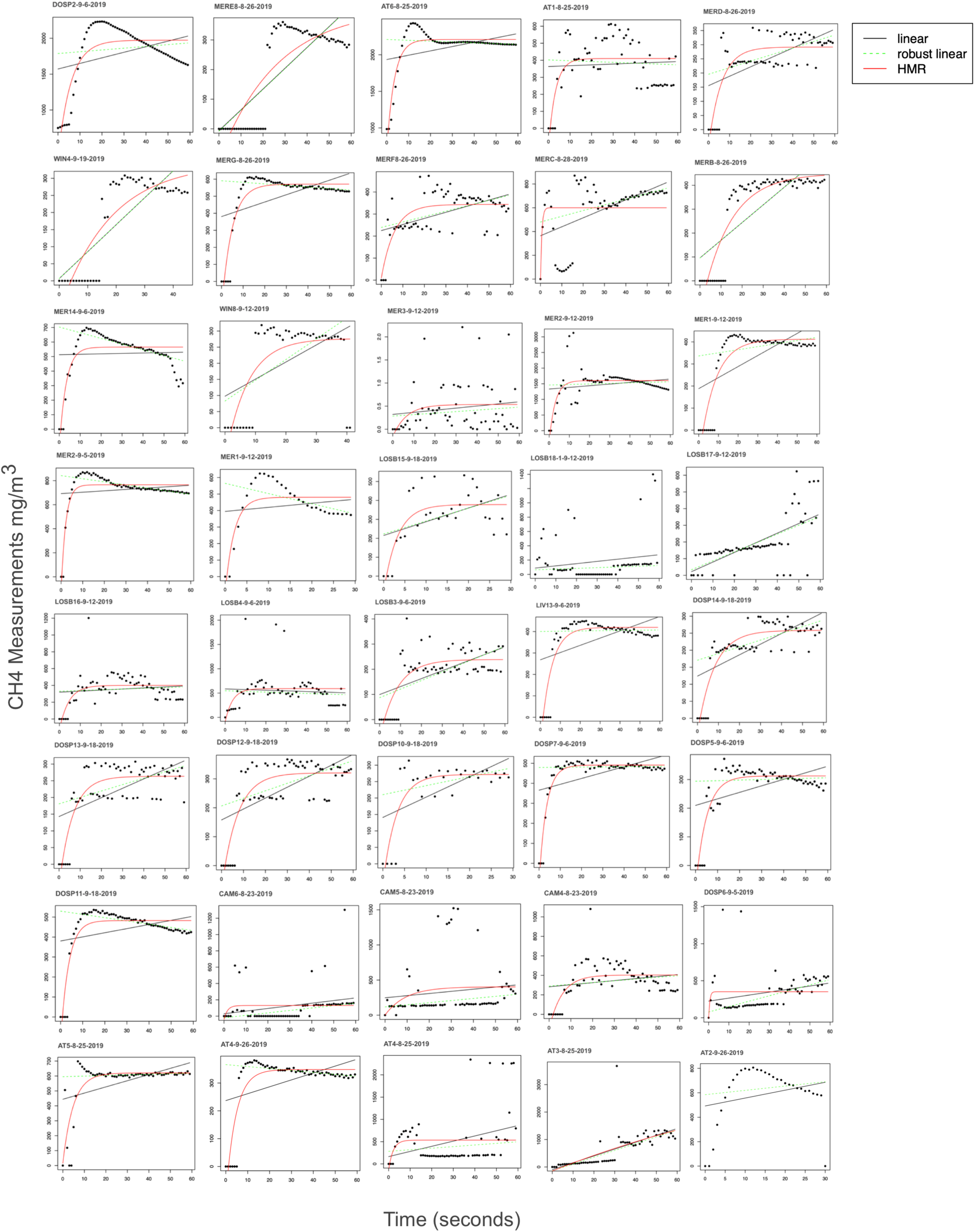
Methane flux calculations for the forty sites that had methane detected by the field gas analyzer. The y-axis is methane measurements in mg/m3and the x-axis is time in seconds. The red line represents Hutchinson-Mosier non-linear modeling, the black line is linear modeling, and the dashed green line represents robust linear modeling. Black dots are the gas measurements at the time point.

### Hotspots

Density and contour plots were generated to map locations of methane and carbon dioxide flux and estimate areas where greenhouse gas flux was likely to occur. (Figure 5). Contour levels represent sample coordinates and greenhouse gas flux data binned to determine regions containing the likeliest density. Density plots estimate both methane and carbon dioxide flux probabilities. The models indicate that carbon dioxide flux is most likely to occur in the region north of Atwater and in an area east of Los Banos, as well as a small region in Merced (Figure 5A,C). Methane flux is likely to occur in Merced city and in the same area east of Los Banos (Figure 5B,D). This geographic region where both methane and carbon dioxide fluxes is are predicted to be highest occur is an unincorporated region named Dos Palos Y located at California State Route 152 and Highway 33 and is approximately 12 miles east of Los Banos and 5 miles north of Dos Palos city in southern Merced County.

**Figure 4.**
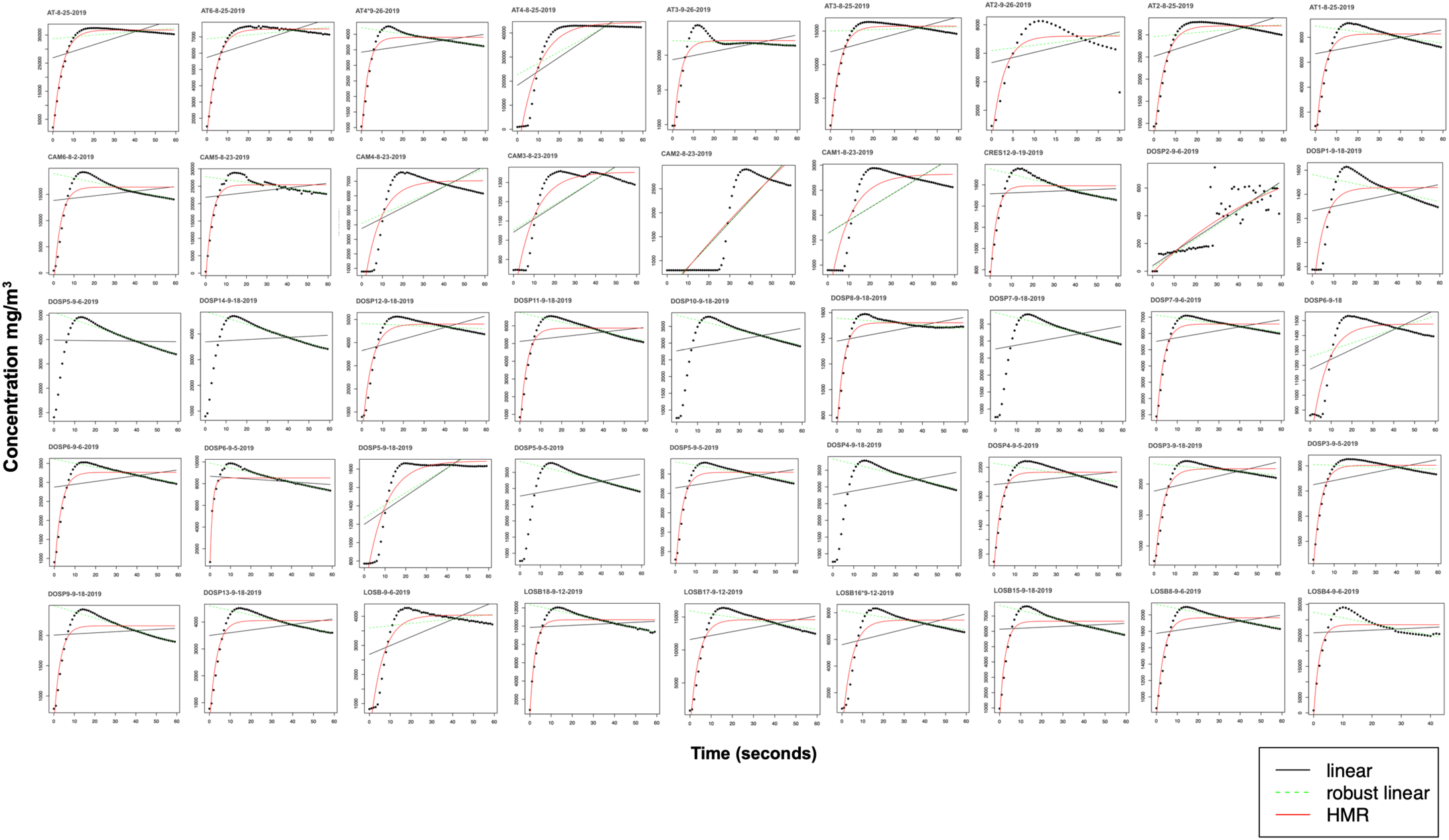
Carbon dioxide flux calculations for the forty-five of the ninety-one sites that had CO2 detected by the field gas analyzer. The y-axis is methane measurements in mg/m3and the x-axis is time in seconds. The red line represents Hutchinson-Mosier non-linear modeling, the black line is linear modeling, and the dashed green line represents robust linear modeling. Black dots are the gas measurements at the time point.

**Figure 5.**
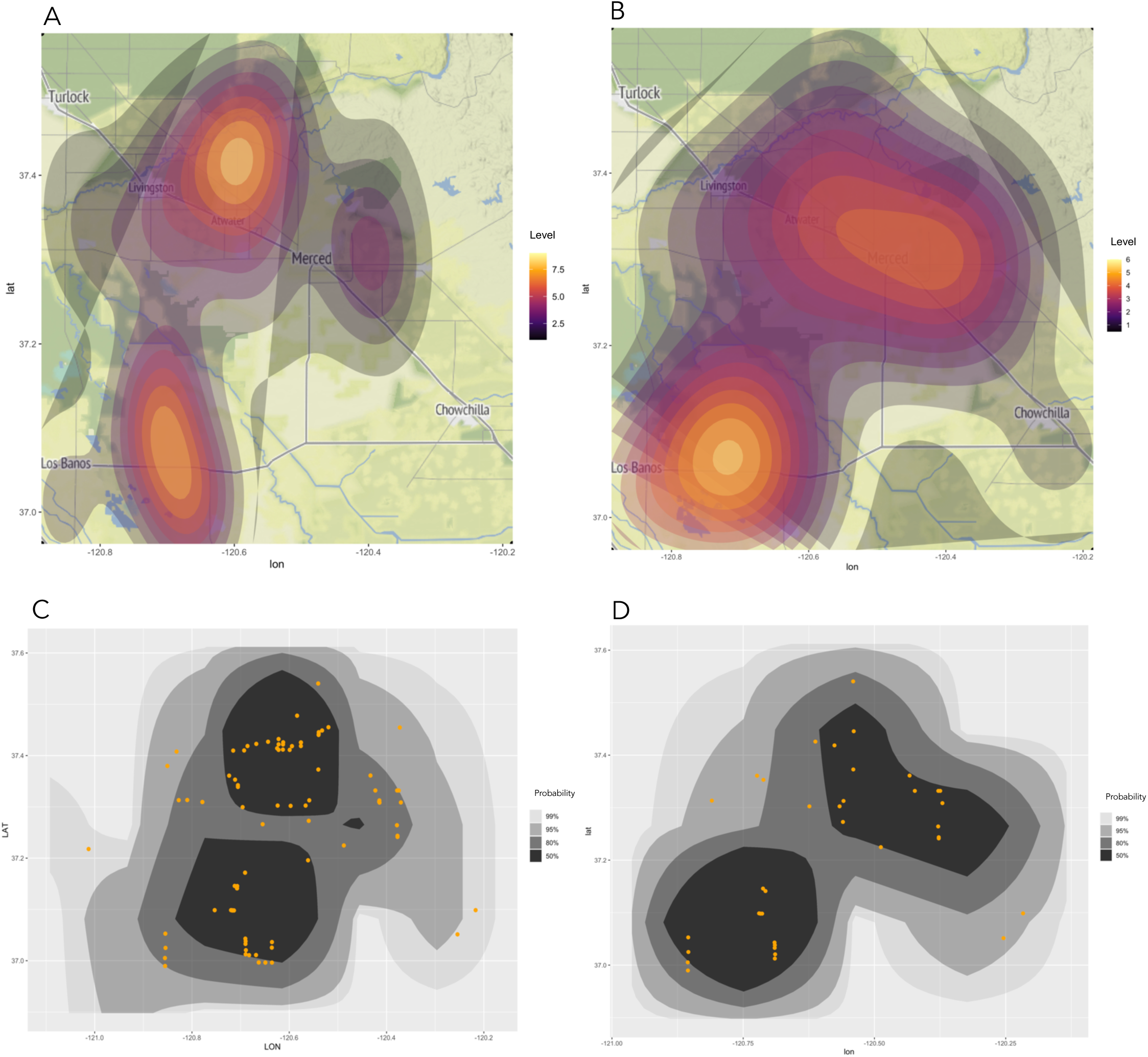
Contour-density plots of methane and carbon dioxide flux. A. Contour-density map of carbon dioxide flux. Levels represents the binning of sampling locations with greenhouse gas flux to determine regions containing the highest density. B. Contour-density map of methane flux. C. Density estimation plot of carbon dioxide flux with each black to gray region indicating probability of highest density; sampling locations with flux are represented by orange dots. D. Density estimation plot of methane flux with each black to gray region indicating probability of highest density.

### Microbial Community Composition and Gas Flux

Taxonomic assignment of metagenome data indicated that Bacteria dominated the irrigation canal sediments, followed by Archaea, Eurkaryota and Viruses. (Figure 6A). From the 38,243 assembled genomes, ten made up 94% of all phyla: Pseudomonadota (45%), Bacillota (20%), Actinomycetota (15%),. Bacteriodota (5%), dsDNA viruses, no RNA stage (4%), Campylobacterota (2%), Euryarchaeota (1%), Spirochaetota (1%), Cyanobacteriota (1%), and Ascomycota (1%). (Figure 6B). All samples contained a similar composition of Archaeal species except for two sediments collected from the Delta-Mendota Aqueduct. There was a small negative correlation between CO_2_ flux and Eukaryotya (correlation: -0.3, p ≤ 0.05) and a small positive correlation between CH_4_ and Spirochaetota (correlation: 0.3, p ≤ 0.05). Planctomycetia was moderately positively correlated with CO_2_ flux (0.4, p ≤ 0.05).

**Figure 6.**
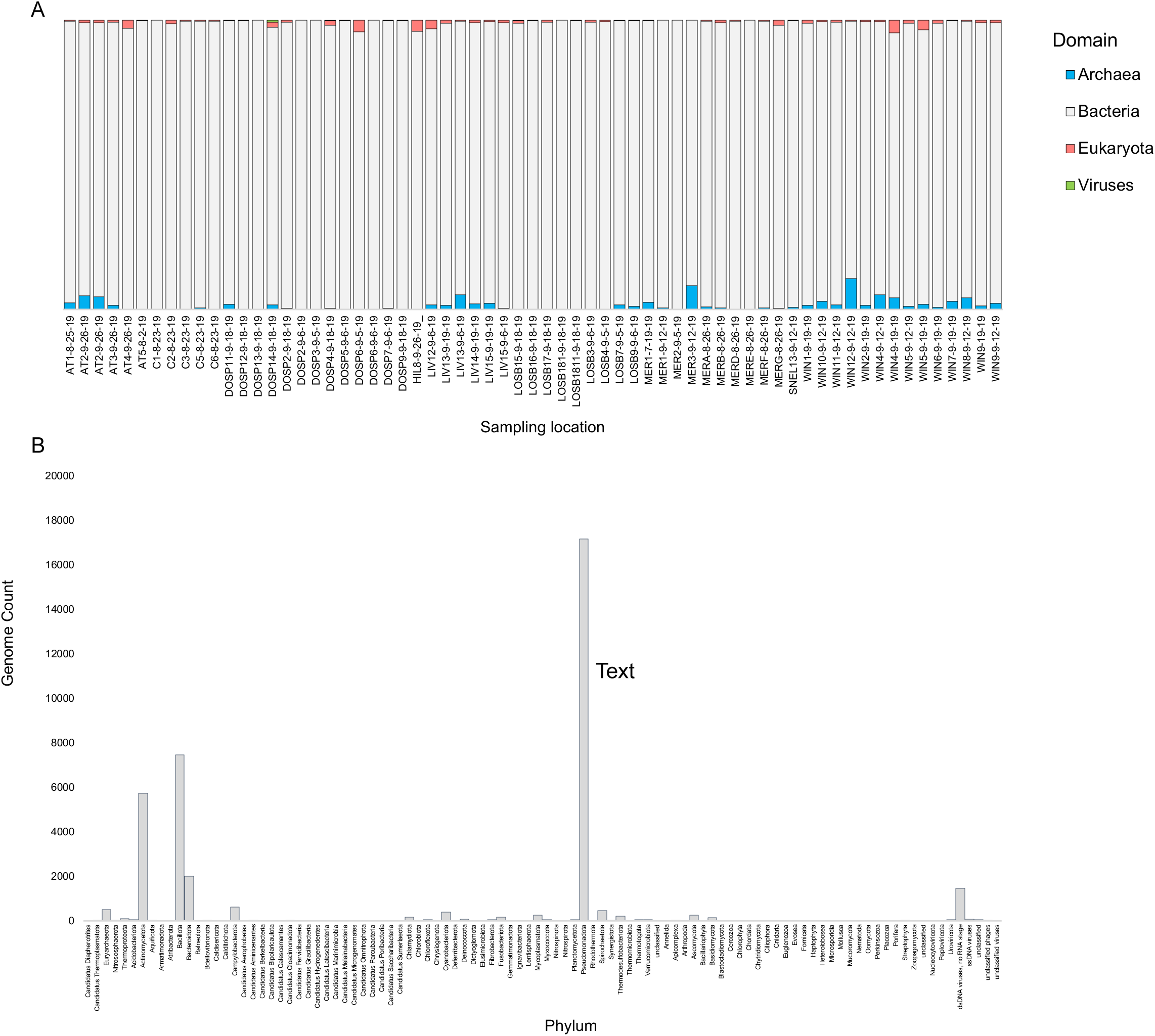
Phylogenetic distribution of 66 sequenced metagenomes. A. Domain distribution across sampling locations. B. Genome counts for 103 phyla.

One hundred-thirty unique species of archaea, 35 of which were methanogens, were identified in the sediment samples. Methanogenic species were identified in all sequenced sediment samples. (Figure 7 (supplemental?)) Samples with a higher percentage relative abundance of methanogenic species were positive for methane flux, while the association was not statistically significant, 63% had methane flux. The most occurring species of methanogens were *Methanoperedens nitroreducens, Methanomassiliicoccus luminyensis, Methanoregula boonei,. Methanothrix soehngenii,* and *Methanoregula formicica*, found in 91% of the sampling locations with successful metagenomic sequence data. The most abundant fFamilies of methanogens were Methanoregula, Methanobacterium, Methanosarcina, Methanothrix, Candidatus Methanoperedens, Methanothermobacter, Methanomassiliicoccus, Methanospirillum, Methanocella, and Methanobrevibacter. (Figure 8, Sup Table 1). Two unique unclassified methanogen fFamilies, within the Methanomicrobiales and Methanosarcsinales, were identified only in one sampling location in Merced (ID: MERE-8-26). Sediment samples LOSB1811-9-18-19 and LOSB1811-9-18-19 were dredged from the Delta-Mendota aqueduct.

**Figure 7.**
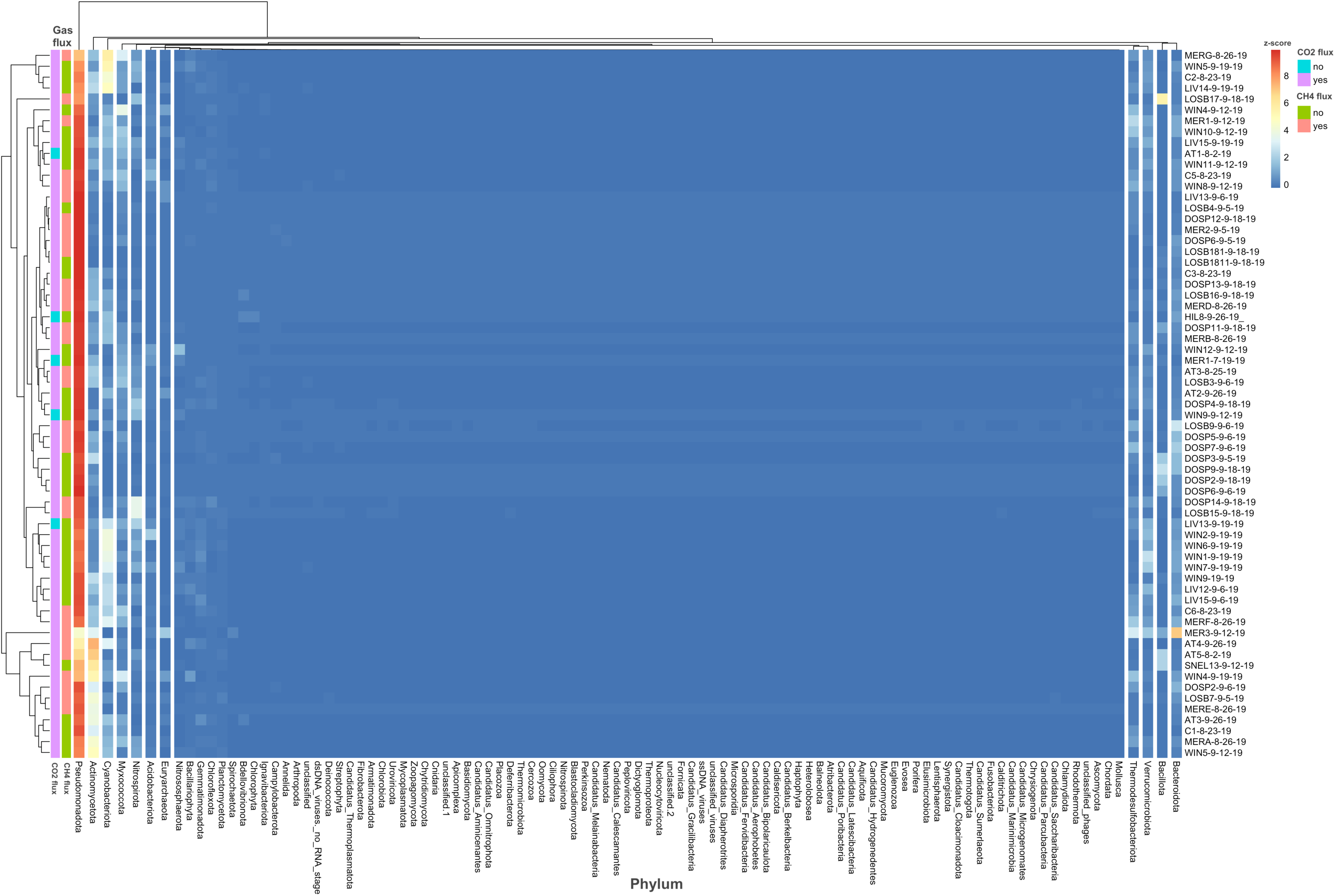
Heatmap of microbial community composition. Sediment samples with are along the right y-axis and gas flux are along the left y-axis. Break inicates clustering of most abundant phyia.

Metagenome assembled genomes (MAGs) were 99% bacterial, including multiple methanotrophic species. Within the MAGs there were 1,544 unique bacteria identified and included two archaeal species, *Methanococcoides burtonii* and *Methanosarcina acetivorans*, and two eukaryotic species, a blight-associated fungus and an algae. Eight sediment samples contained 15 unique methanotrophic methane oxidizing bacterial genera. MAGs for ammonia oxidizing bacteria. *Nitrosococcus, Nitrosomonas,* and,*Nitrosospira* were also identified from sediment samples.

### Microbial Functional Profiles

Metabolic pathways associated with methanogenesis were identified in the metagenomic data. We generated functional profiles using MetaCyc pathways (Figure 9). Methanogenesis from acetate, trimethylalamine, and H_2_ and CO_2_ were the most abundant across all samples. Methanogenic pathways starting from acetate and trimethylamine were the most abundant in sediment samples that had both methane and carbon dioxide flux. Methanogenesis pathways starting from methylthiopropanoate, methanethiol, and dimethylsulfide were moderately positively correlated with carbon dioxide flux (0.4, p<= 0.005). MetaCyc pathways for methane metabolism starting from tetramethylammonium (MetaCyc: PWY-5261) and glycine betaine (MetaCyc:PWY-8107) were not identified in any of the samples.

**Figure 8.**
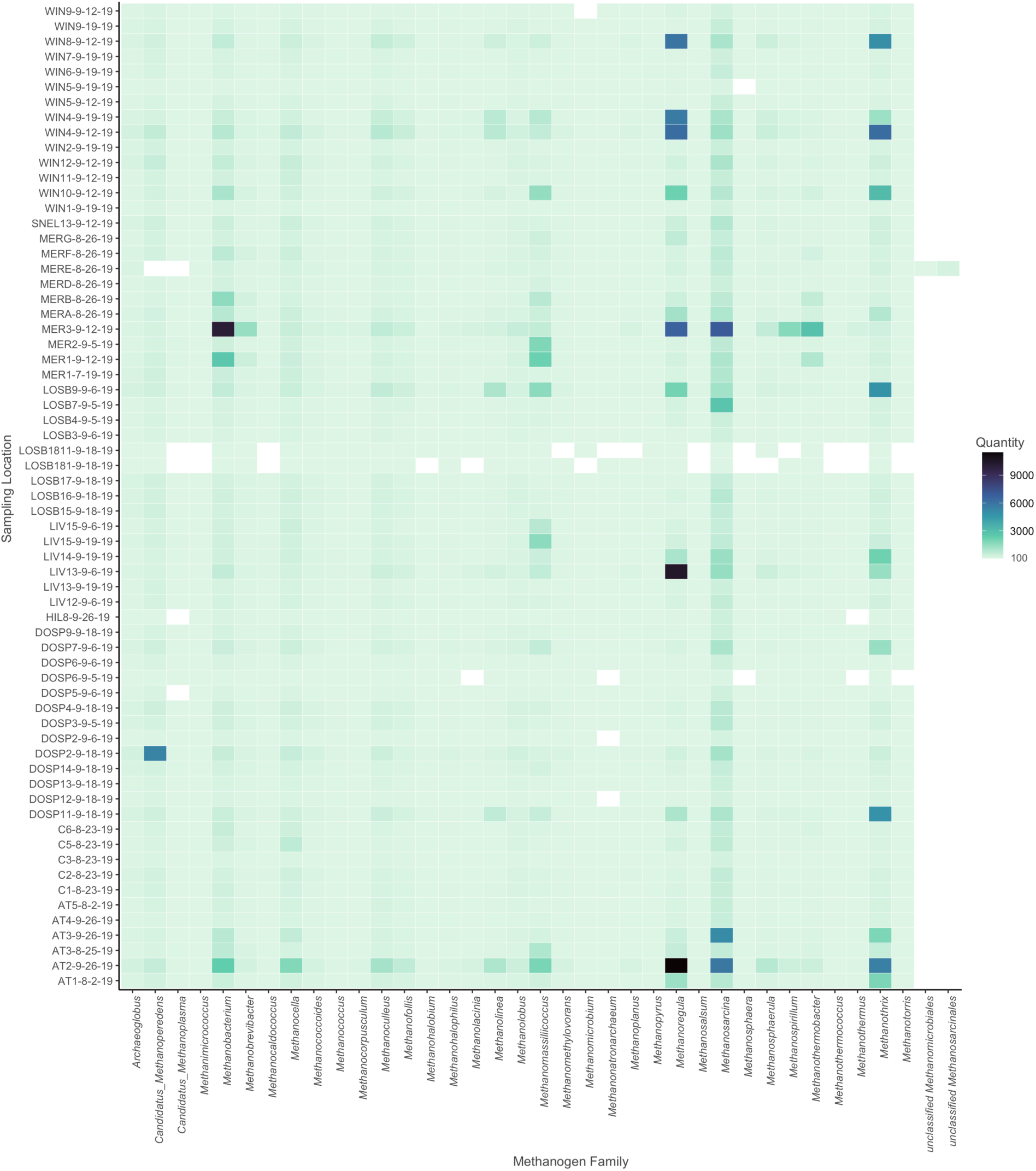
All methanogenic genera per sampling site as identified by metagenomic sequencing.

**Figure 9.**
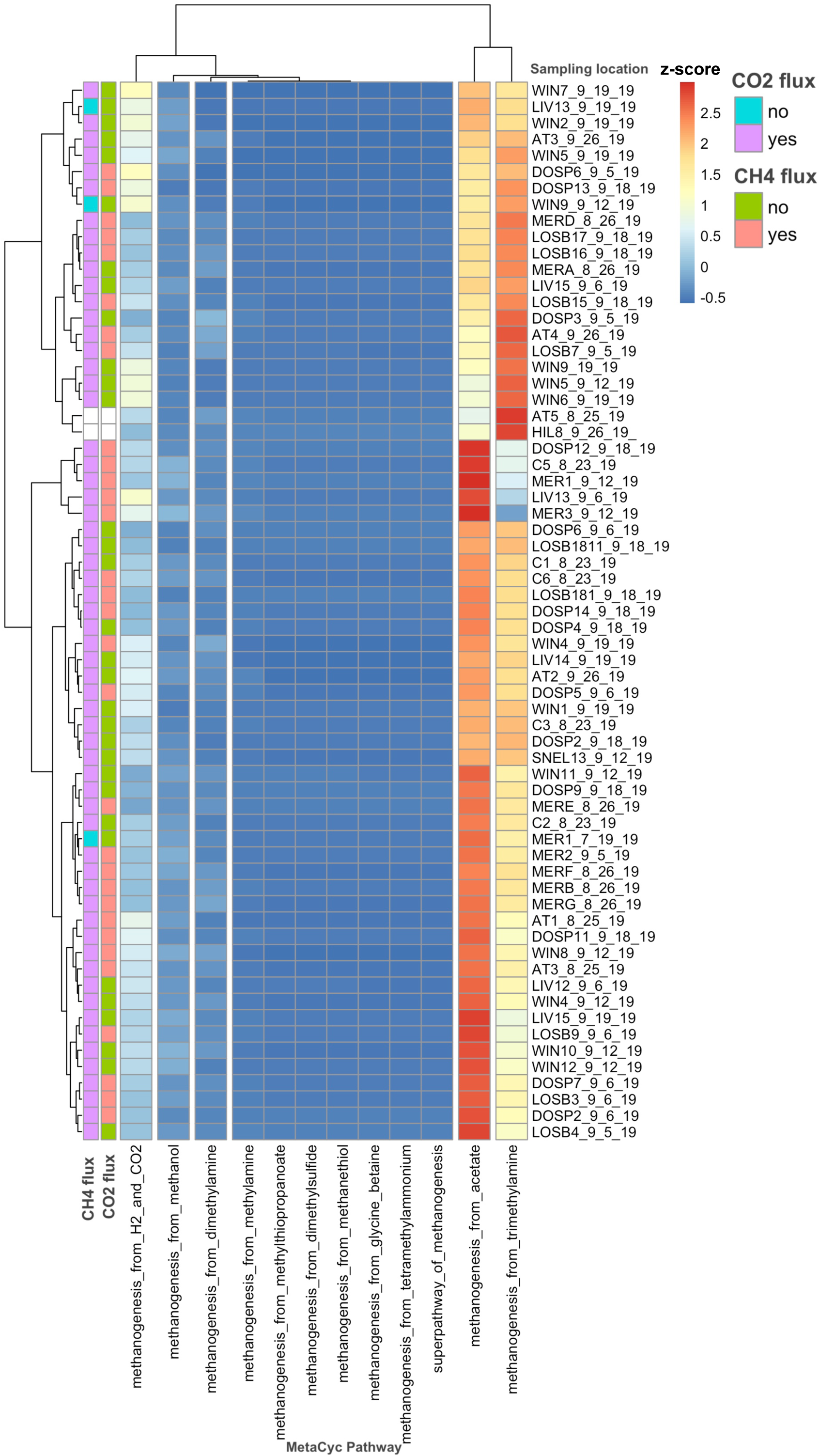
Microbial functional pathways identified in metagenomic sequencing data. Sediment samples are on the right y-axis and gas flux is on the left y-axis. Break occur for grouping of most abundant MetaCys pathway results.

**Figure 10.**
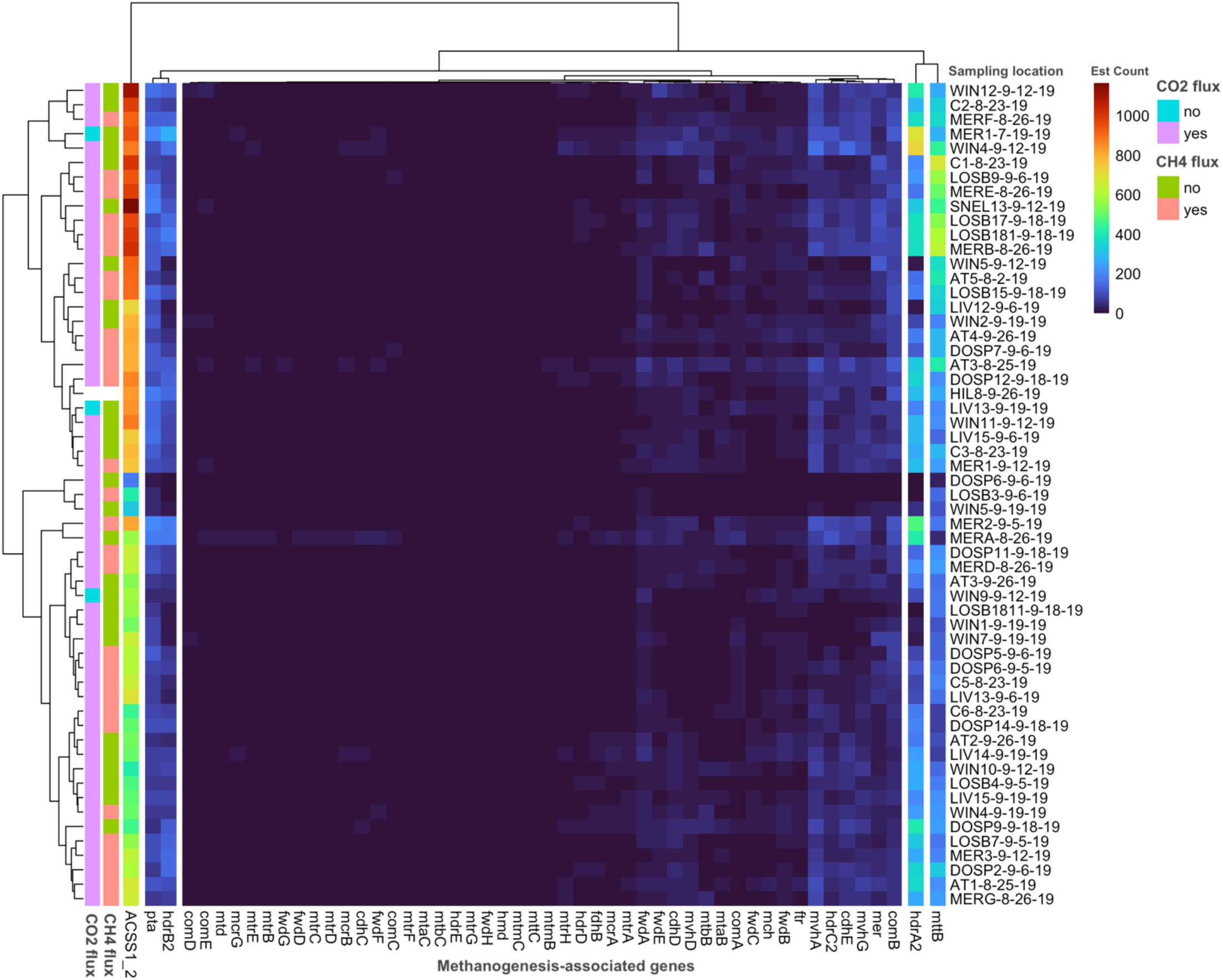
Methanogenesis-associated genes from metagenomic sequencing per sampling site.

MAG genera associated with methane metabolism and methane oxidizing bacterial groups were both positively and negatively correlated with methane and carbon dioxide emission. Methane was moderately negatively correlated with *Methylibium* and *Methyloversatialis* (-0.4, p<=0.5). Methane was strongly positively correlated with *Methylomicrobium* (0.7, p=0.05) and moderately correlated with *Methylocaldum*(0.5, p<=0.5). Carbon dioxide was strongly positively correlated with *Methylobacillus* (0.8, p<= 0.05) and moderately correlated with *Methylosarcina* and *Methylocaldum* (0.4, p<=0.5). Carbon dioxide was moderately negatively correlated with *Methyloversatilis* (-0.5, p<=0.5) and *Methylibium* (-0.4, p<=0.5).

Several enzymatic genes associated with methanogenesis were identified in all sequenced sediment samples. (Figure X). Five enzyme genes, acetyl-CoA synthetase (ACSS1_2), hdrA2, hdrB2, trimethylamine-corrinoid protein Co-methyltransferase (mttb), and phosphate acetyltransferase (pta) were highly abundant and found in samples that had were both positive and negative for methane and carbon dioxide flux. Methane flux was moderately positively correlated with methanogenesis-associated enzymes 5,10-methenyltetrahydromethanopterin hydrogenase (Hmd, 0.5, p<=0.05), methanol 5-hydroxybenzimidazolylcobamide

Co-methyltransferase (mtaB), methanol corrinoid protein (mtaC), 2-phosphosulfolactate phosphatase (comB) and L-2-hydroxycarboxylate dehydrogenase (NAD+) (comC) (0.3, p<=0.05). (Sup figure X) Genes for superoxide dismutases were identified in the MAGs (Sup table) but were not identified in Archaea. However, standard assembled genomes for *Methanosarcina barkeri* contained superoxide dismutase, Cu-Zn family SOD1. Kegg orthology identifiers for enzymatic genes belonging to methanogenic pathways were also quantified. (Figure 10)) The most abundant KO-associated gene was KO1985, acs, acetyl-CoA synthetase from KEGG Module M00357 (acetate to methane). The second most abundant was KO3388, hdrA2 heterosulfide reductase subunit A3 from KEGGegg Modules for methanogenesis, M00356 (methanol), M0357 (acetate), M00563 (methylamine/ dimethylamine/ trimethylamine), and M00567 (carbon dioxide).

## Discussion

Engineered terrestrial aquatic systems’ contribution of greenhouse gas to the atmosphere is important to quantify when developing strategies to mitigate global climate change. Irrigation canals for large-scale industrial agriculture make up thousands of miles of these engineered systems and have not been a focus of research. It is unknown if the irrigation canals in the agriculture-heavy Central Valley of California are emitting more than one type of greenhouse gas. Due to their acquisition of fertilizer-rich soil, livestock wastes, and other agricultural byproduct runoff that con contribute to the establishment and metabolic support of microorganismal communities, these canals have strong potential to contribute to greenhouse gases to the atmosphere.

This study addresses whether engineered aquatic systems are significant sources of greenhouse gas emissions. Methane and carbon dioxide fluxes measured in the field were paired with microbial community characterization via culture-free metagenomic sequencing of irrigation canal sediments in the Central Valley of California. Measurements at one hundred locations showed high levels of carbon dioxide flux at nearly all sites and methane flux at close to a third of sites, indicating that the canals are releasing greenhouse gases. Methanogenic and methanotrophic species were present in all the sequenced sediment samples and moderately correlated with methane and carbon dioxide production at their sampling locations. This is consistent with the emerging recognition of the impact that irrigation canals and other engineered terrestrial aquatic systems can have on global climate change. Peacock et al 2021, urged inclusion of ‘artificial’ drainage ditches and irrigation canals in global emission data and demonstrated that methane emission occurs from drainage ditches.^30^

This work contributes to the growing body of studies demonstrating that methane and carbon dioxide flux can be associated with irrigation systems ^31,32,33,34^ The correlation we found between carbon dioxide flux and Planctomycetota reflects the known role of this phylum in global carbon cycling.^35^ While most methanogens are unable to survive oxygen-rich environments such as aerated water in systems with constant flow and movement, there are several oxygen-tolerant genera including *Methanobrevibacter* and *Methanosarcina*, both of which were identified in the sequenced sediments.^36^ *Methanosarcina barkeri* contained superoxide dismutase, SOD1, and this enzyme could aid methanogenesis in an oxic environment.^37^ Presence of ammonia oxidizing bacteria in the canal sediments indicate that methane flux may be countered by the consumption of methane and could be the explanation for the carbon dioxide flux found at 91% of the sampled locations.^38,39,40^ However, multiple studies suggest that methanotrophs may mitigate climate change.^41,42,43^

This study was conducted in the summer months and addresses two main aspects of greenhouse gas emission methodology; gas flux and microbial community profiling. High ambient temperatures during field testing and sampling (99 – 115°F) caused equipment failure and excluded sediment temperature, pH and water flow rate data. However, while we initially hypothesized flow rate to have a large impact on the presence of sediment, plant and microorganisms, the fastest moving system, the aqueduct, and other rapidly moving sections of the canals were rich in plant and animal life along with sediment at the sides and basins. Processing the entire sediment collection for metagenomic sequencing would provide a complete dataset to more robustly supported predictions. The region hotspot highlighted by the contour-density plots, Dos Palos Y, one we were not able to access fully, is the site of a very large livestock auction business. Cattle are the main animal housed on this site. The addition of nitrate and ammonia as fertilizer to crops in the Central Valley and subsequent runoff into the irrigation canals likely provides alternative electron acceptors allowing methanotrophs to process methane and release carbon dioxide.^44^

*Methanobrevibacter* and *Methanosarcina*, both of which were identified in the sequenced sediments.^36^*Methanosarcina barkeri* contained superoxide dismutase, SOD1, and this enzyme could aid methanogenesis in an oxic environment.^37^ Presence of ammonia oxidizing bacteria in the canal sediments indicate that methane flux may be countered by the consumption of methane and could be the explanation for the carbon dioxide flux found at 91% of the sampled locations.^38,39,40^ However, multiple studies suggest that methanotrophs may mitigate climate change.^41,42,43^

## Conclusion

Accumulated sediments in the Central Valley irrigation canal and aqueduct system create unique ecological niches that harbor complex microbial communities that impact ecosystem nutrient cycling through methane and carbon dioxide production.

## Data

Data can be accessed at JGI IMG. Project information is maintained at: https://gold.jgi.doe.gov/study?id=Gs0145339

---

**Table 3.**
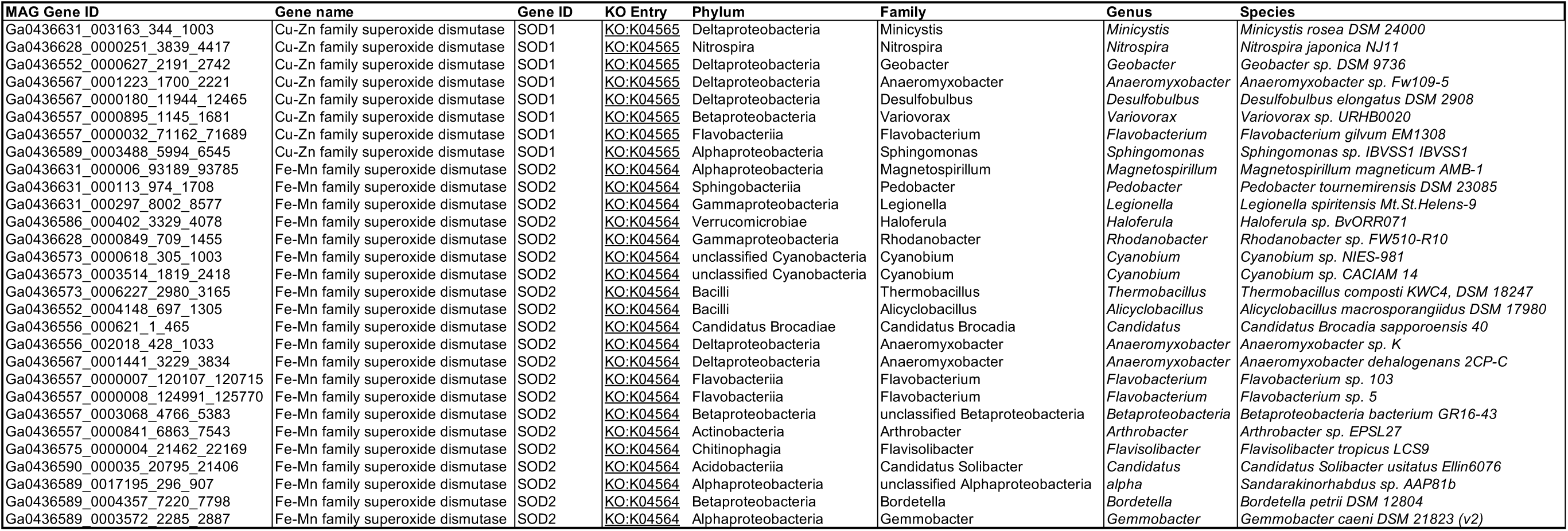
Distribution of metagenome assembled genomes.

**Sup Figure 1.**
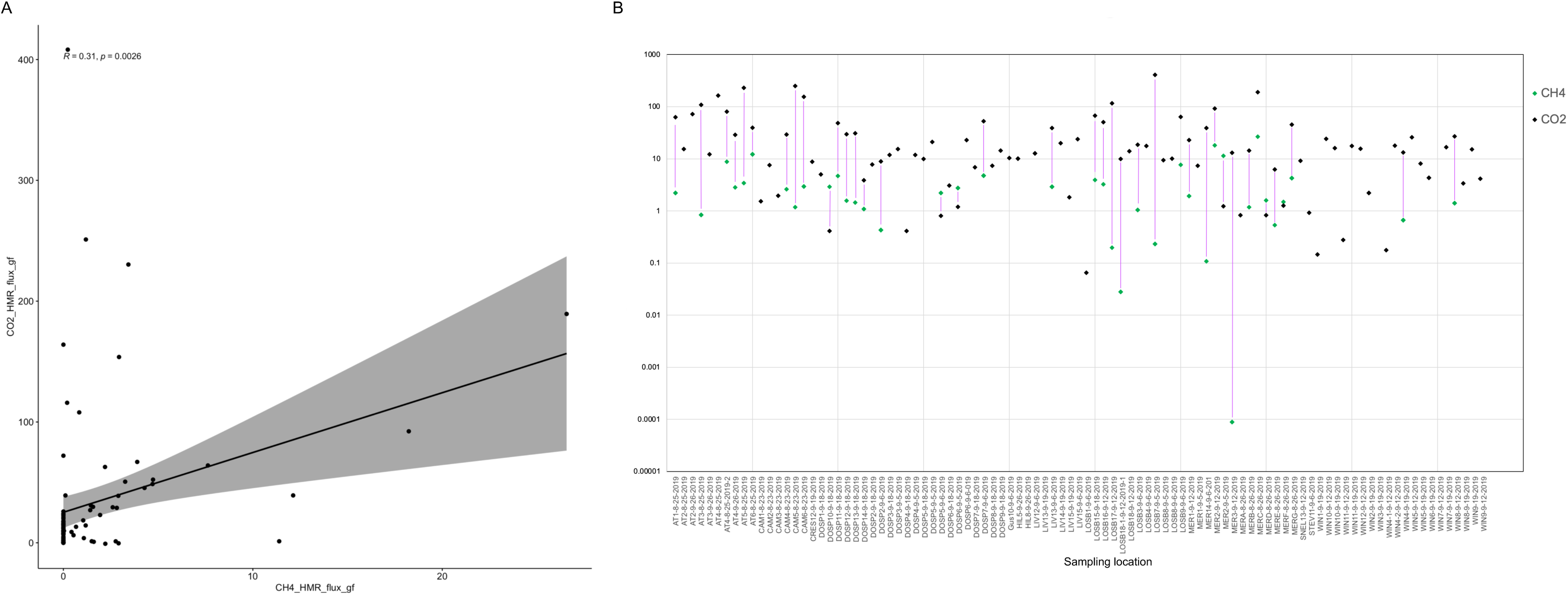
A. Correlation plot between carbon dioxide flux and methane flux. There is a moderate positive correlation between the two greenhouse gas fluxes. (p ≤ 0.005) B. Methane and carbon dioxide flux co-occurrence, log transformed. Black diamonds represent carbon dioxide and green diamonds represent methane flux. The connecting pink lines indicate the co-occurrence of both greenhouse gas flux measured per sampling location; length does not represent any additional information.

**Figure X.**
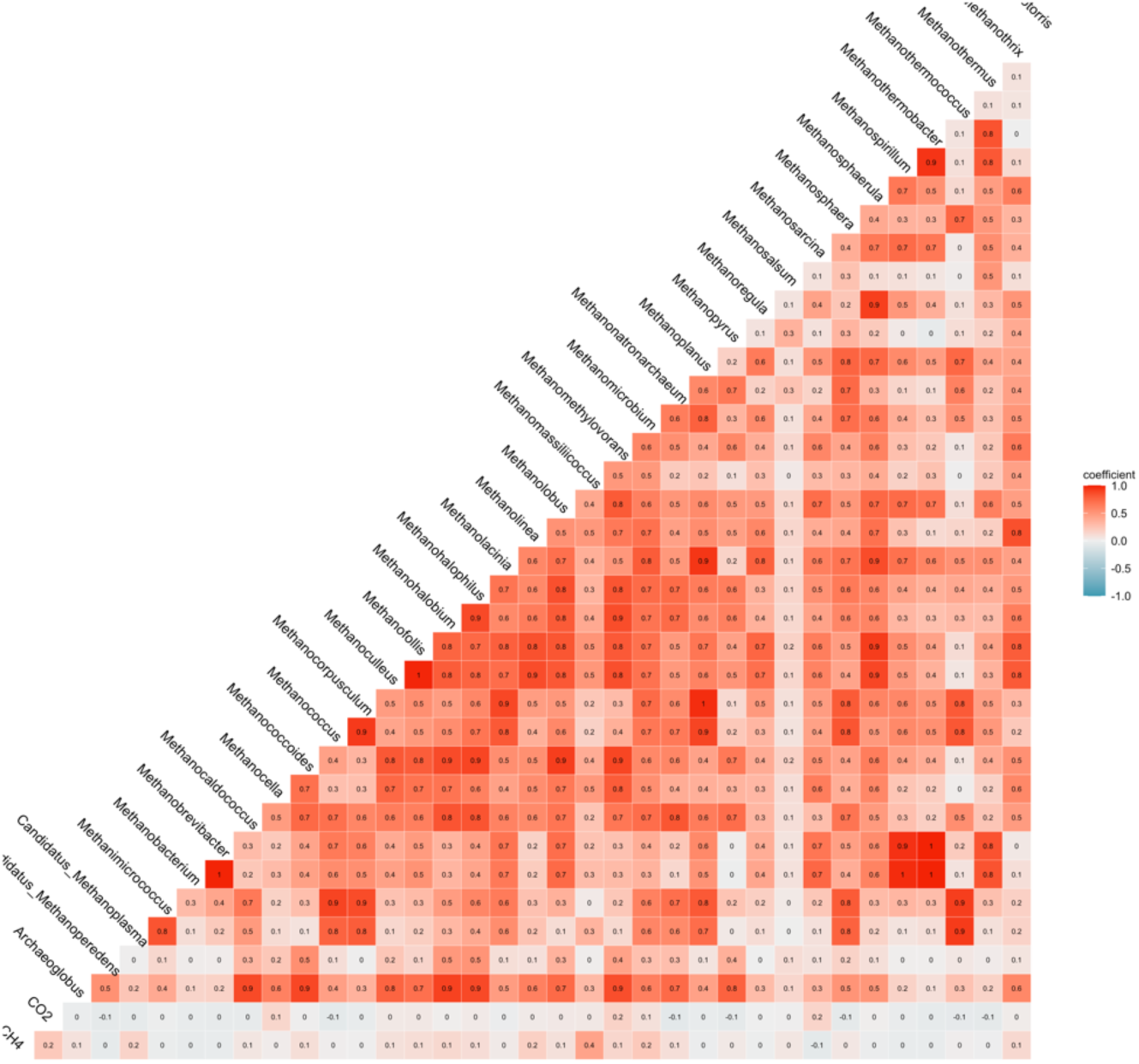
Correlation matrix for methanogens identified by metagenomic sequencing and methane
adn carbon dioxide flux. Moderate positive correlation between CH4 flux and the identification of Methanolobus genera (0.4). There were many instances of high correlation between methanogenic genera themselves. See table

**Figure X.**
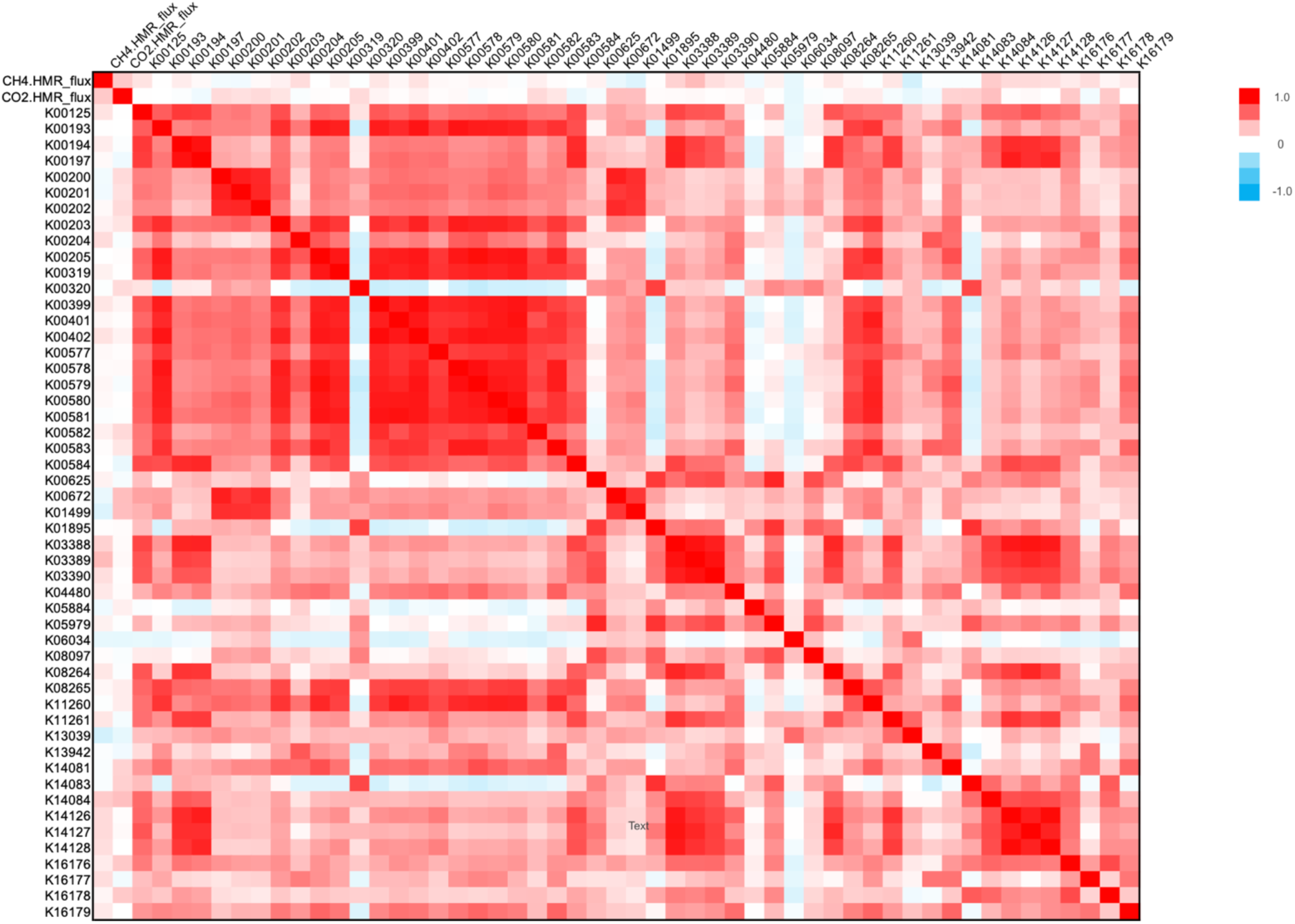
Correlation matrix for methane and carbon dioxide flux values and quantification of methanogenesis associated pathway genes, represented here by Kegg Orthology identifiers.

**Figure X.**
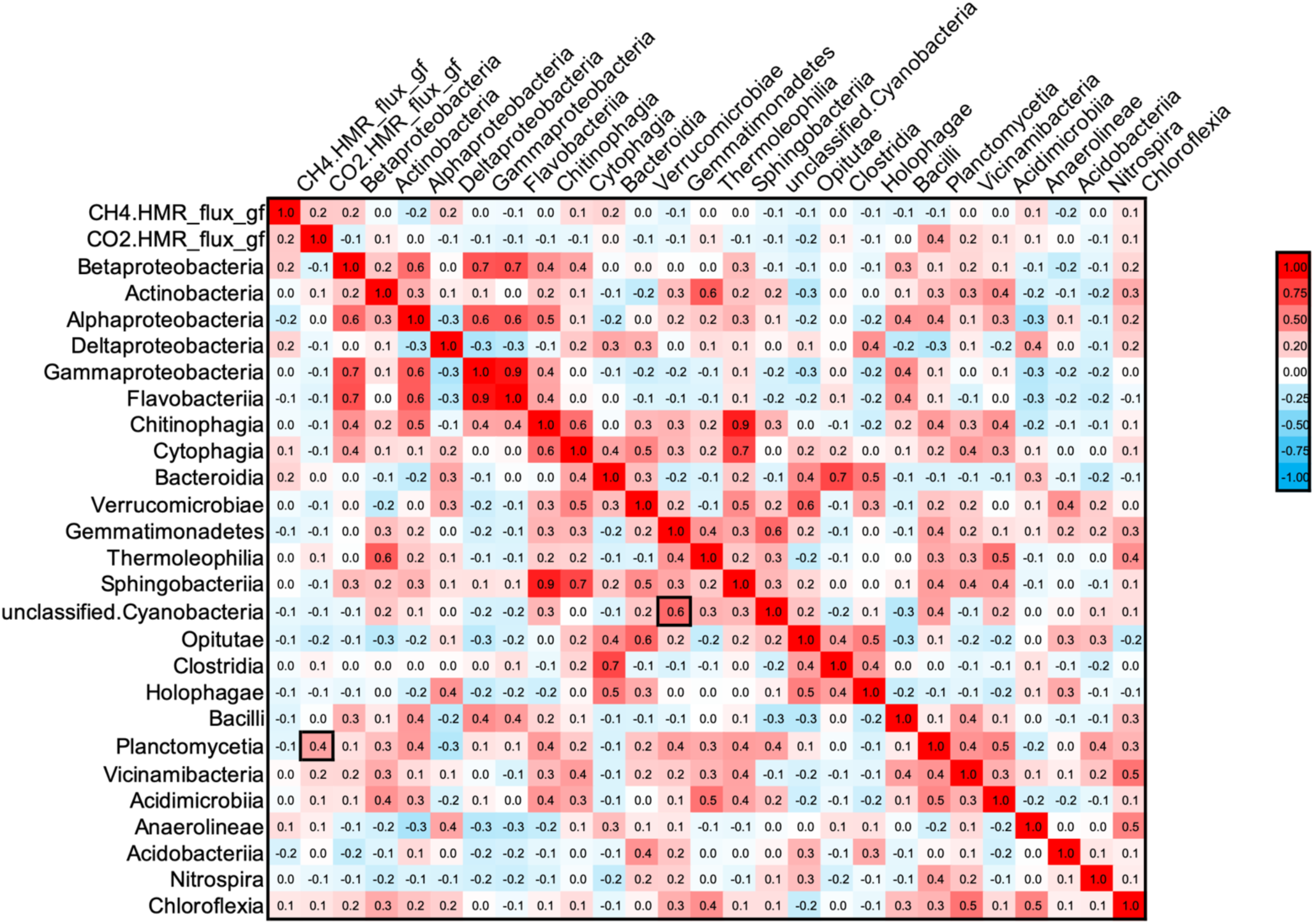
Correlation plot between top 25 most occurring bacteria phyla and methane and carbon dioxide flux calculations. Dark boxes in the interior plot indicate two points of interest.

**Figure X.**
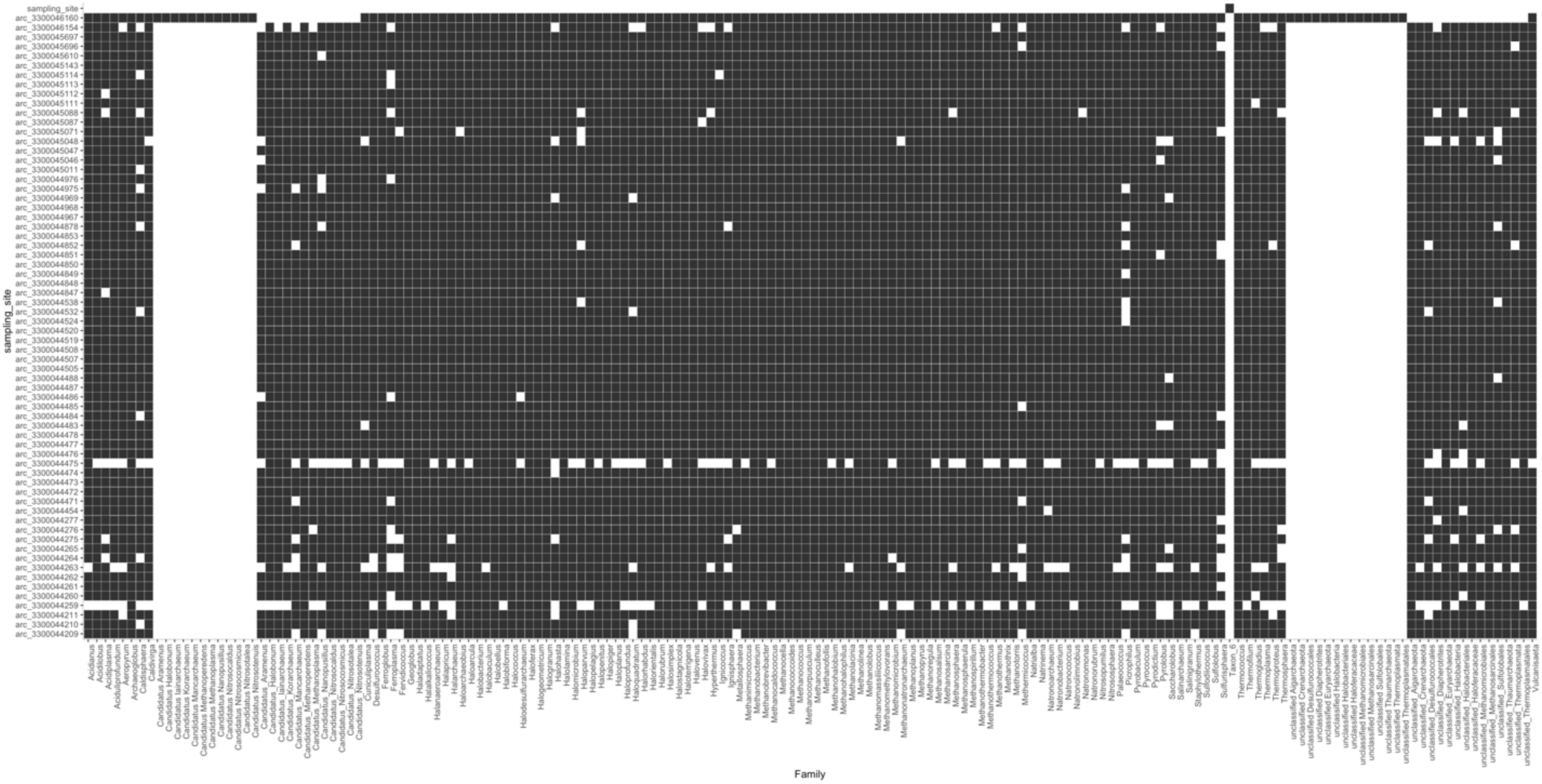
Presence absence matrix for all archaeal genera identified by metagenomic sequencing by sampling location. Black indicates presence of genera. What was special about the top site? 3300046160/MERE-8-26-19

**Figure X.**
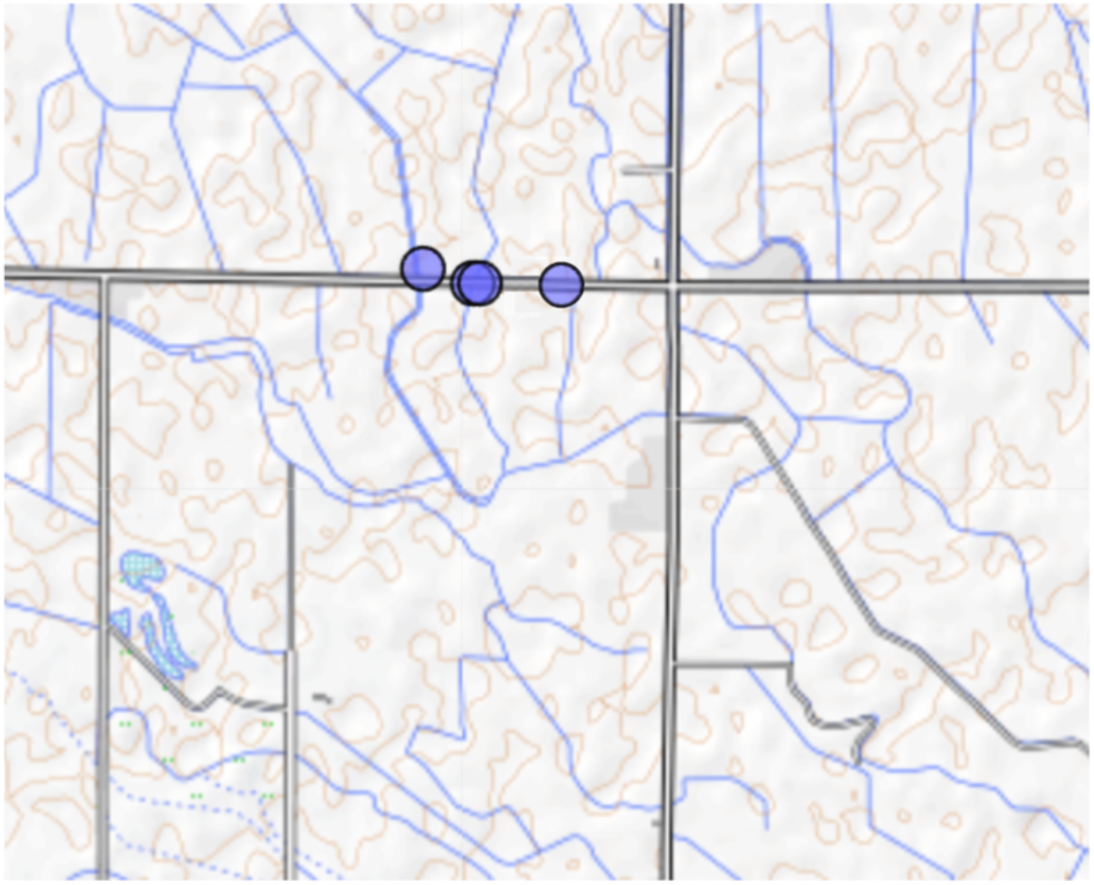
Zoomed in image of overlapping sampling location with plotted points indicating that four different irrigation canals were sampled along the same road. There are two overlapping points in the center indicating one irrigation canal that ran northsouth and one perpendicular to it that ran parallel along the road.

**Figure X.**
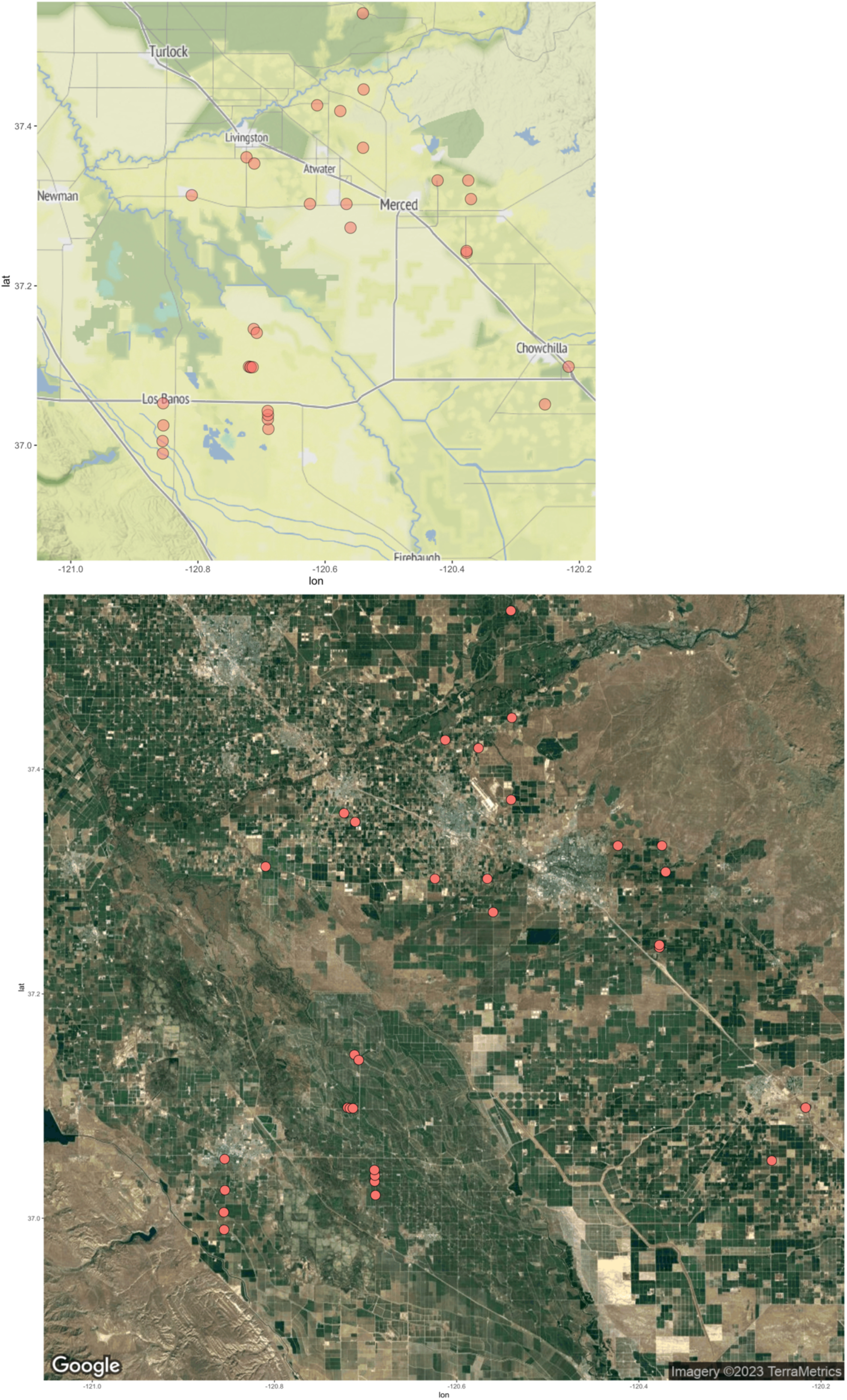
Methane flux locations in the central valley. Overlapping points indicate separate waterway systems. A. lskdlks lksdklsd B. Satellite view indicates agricultural layout sdlksdfml?????

**Sup Figure X.**
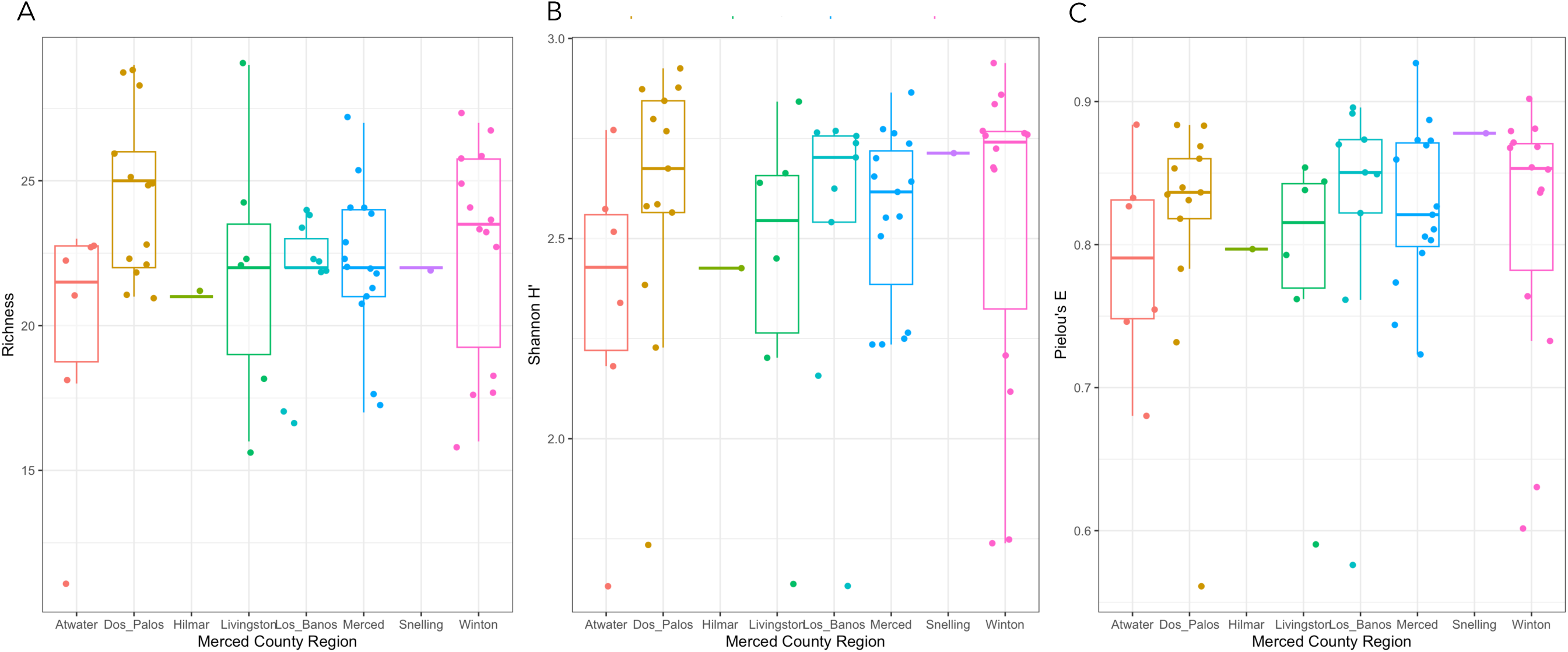
Alpha diversity calculations for methanogens identified by metagenomic sequencing by Merced County Region. A. Species richness. B. Shannon index. C. Pielou’s evenness calculation.

**Sup Figure X.**
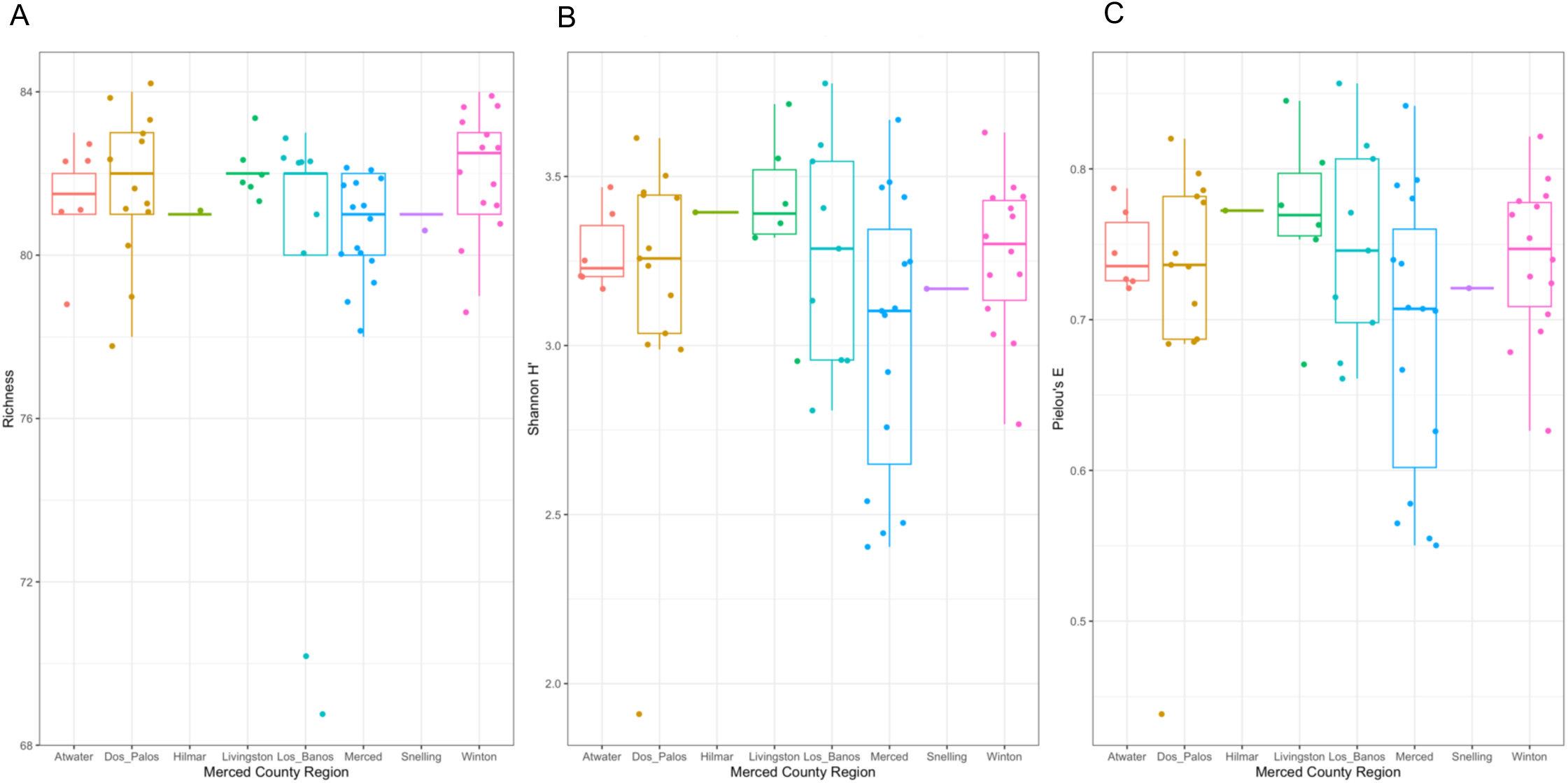
Alpha diversity calculations for bacterial phyla identified by metagenomic sequencing by Merced County Region. A. Species richness. B. Shannon index. C. Pielou’;s evenness calculation.

**Sup Figure X.**
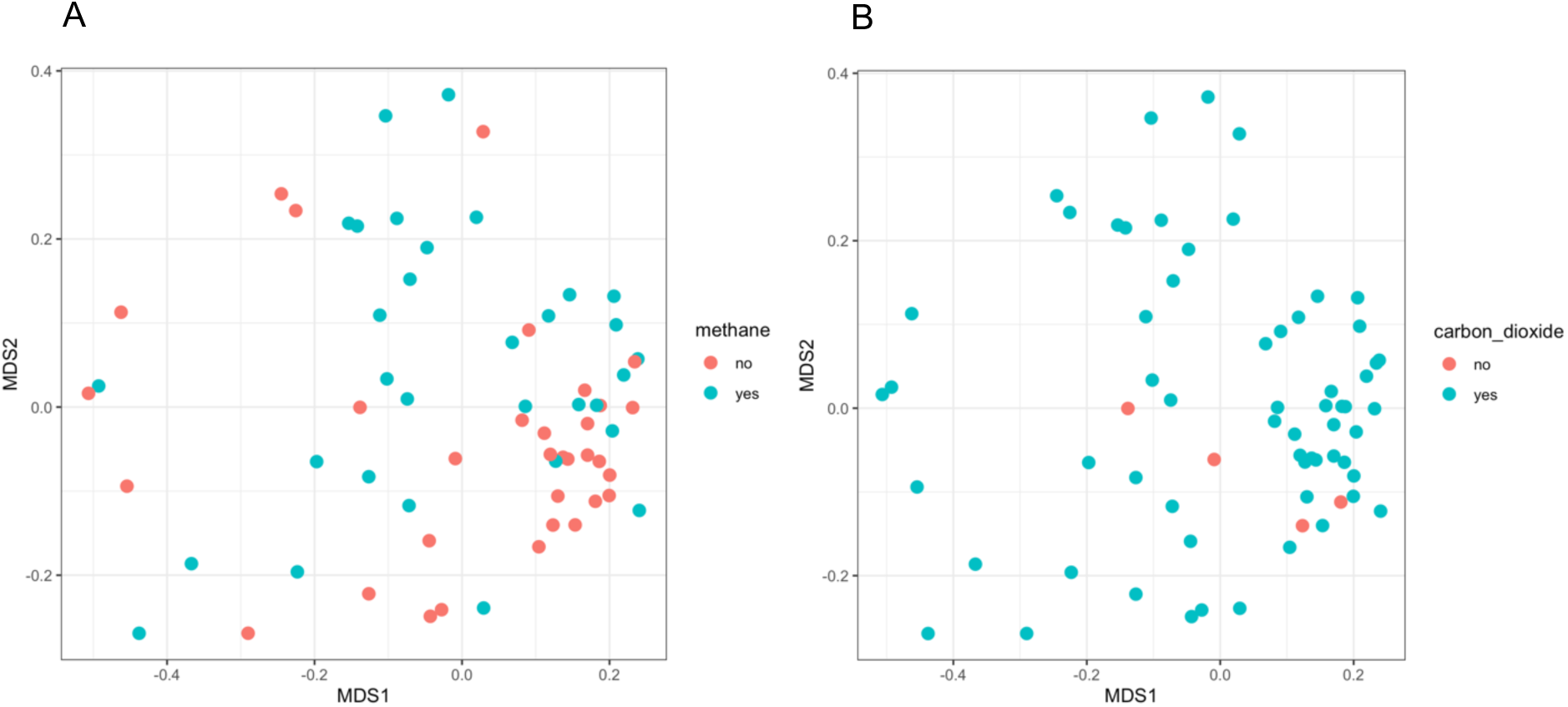
MSDS plots of sampling locations with identified bacterial phyla from metagenomic sequencing. A. MSDS ordinal plot by methane flux. B. MSDS ordinal plot by carbon dioxide flux. Neither ordination explains clustering.

**Figure X.**
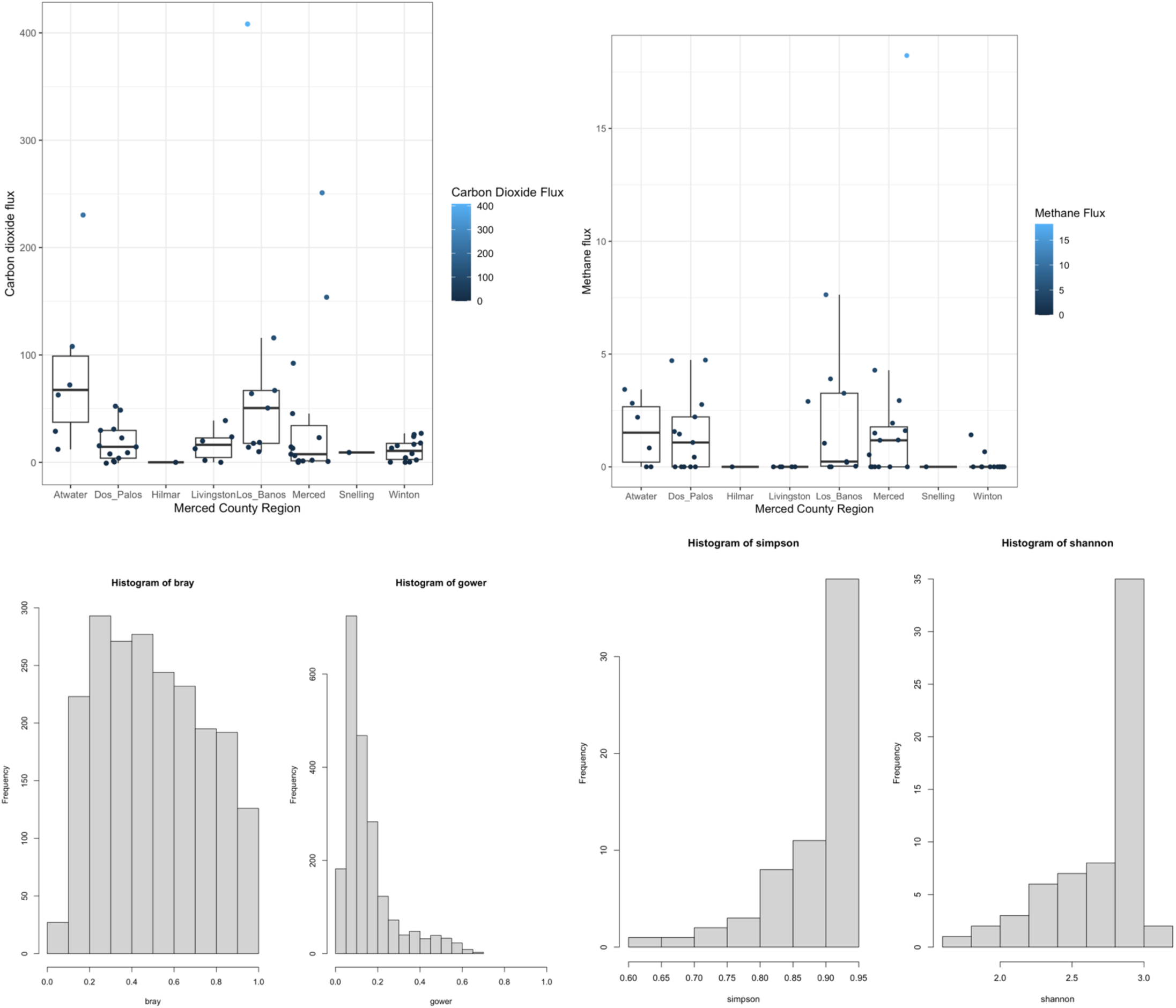
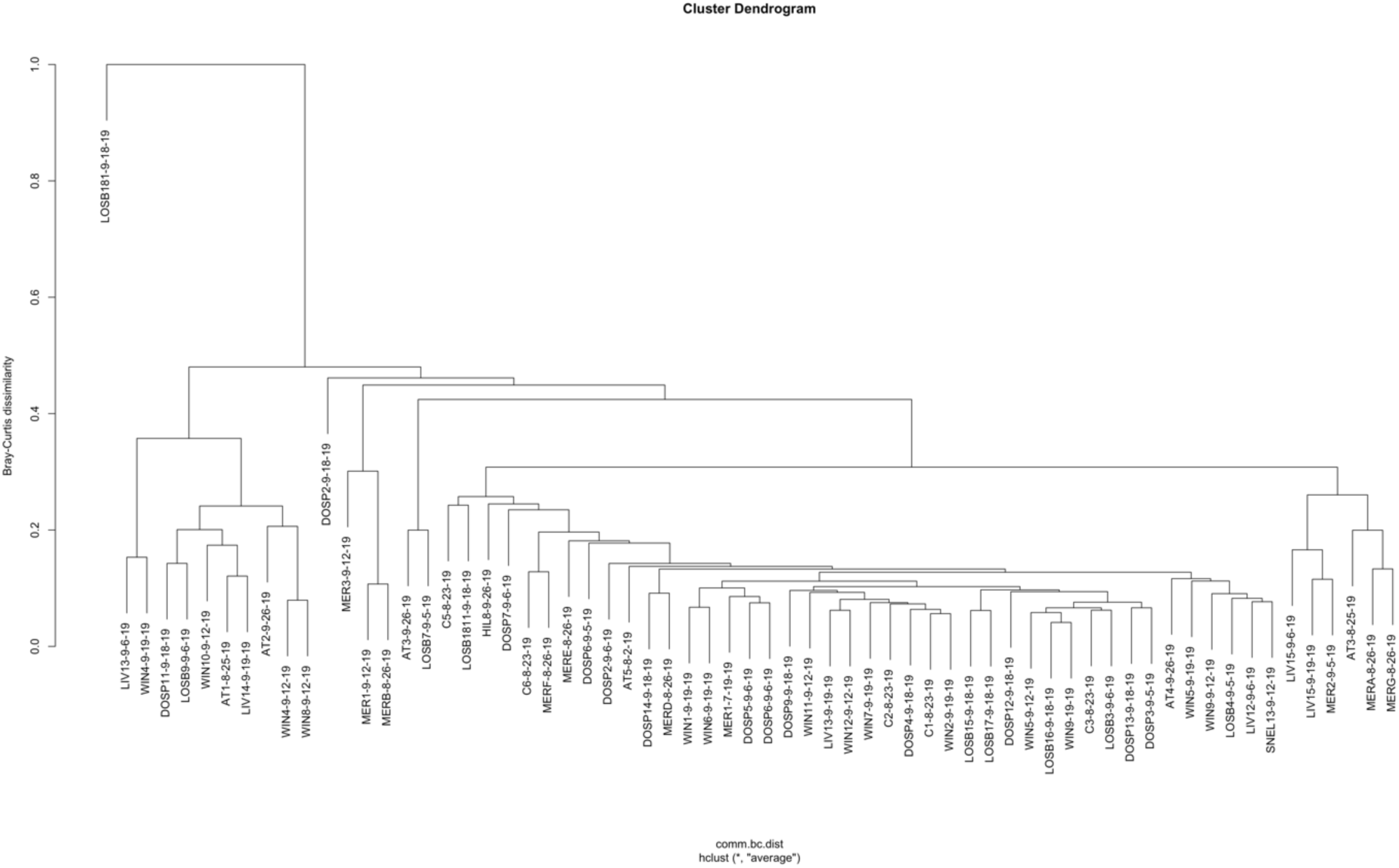
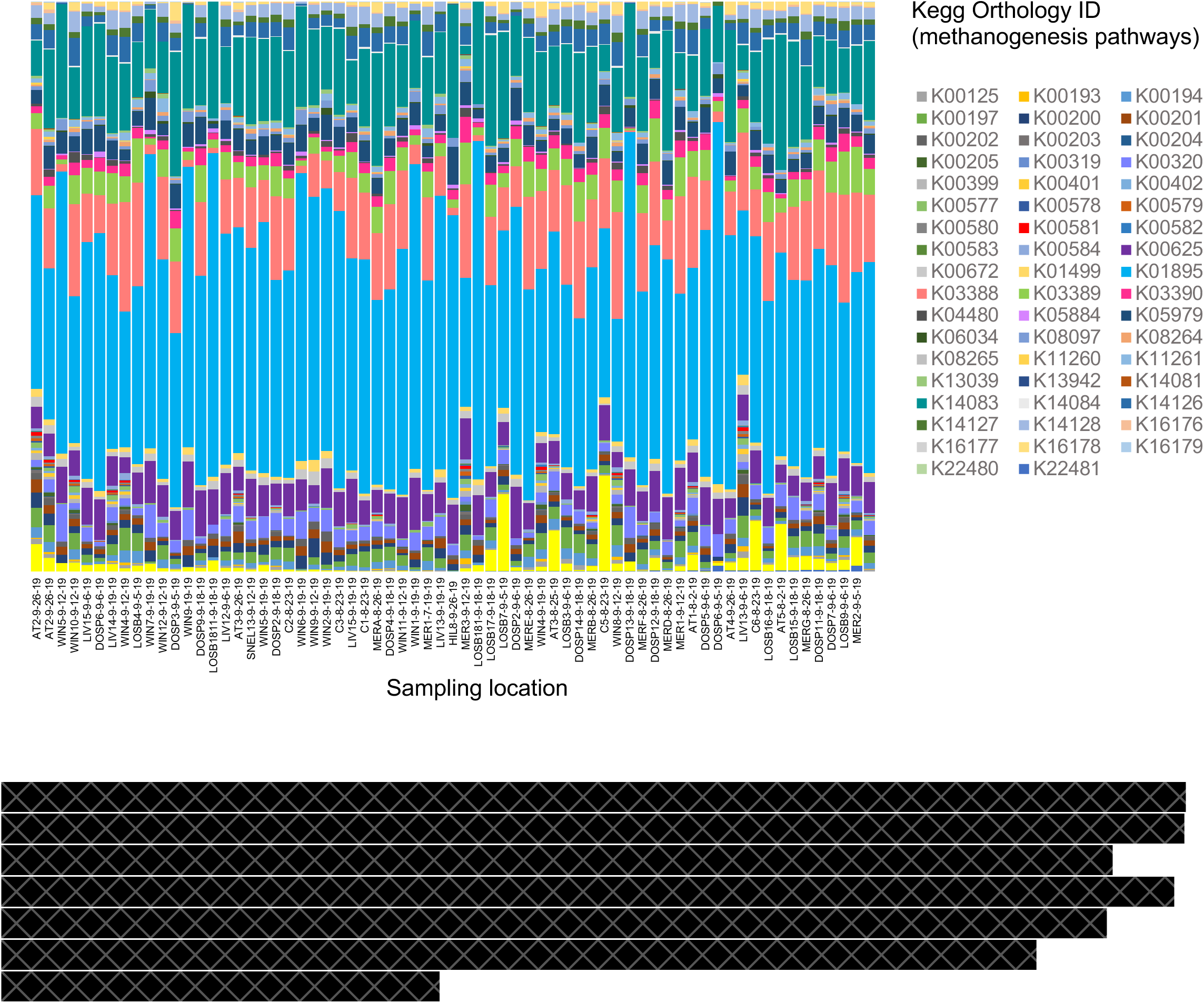
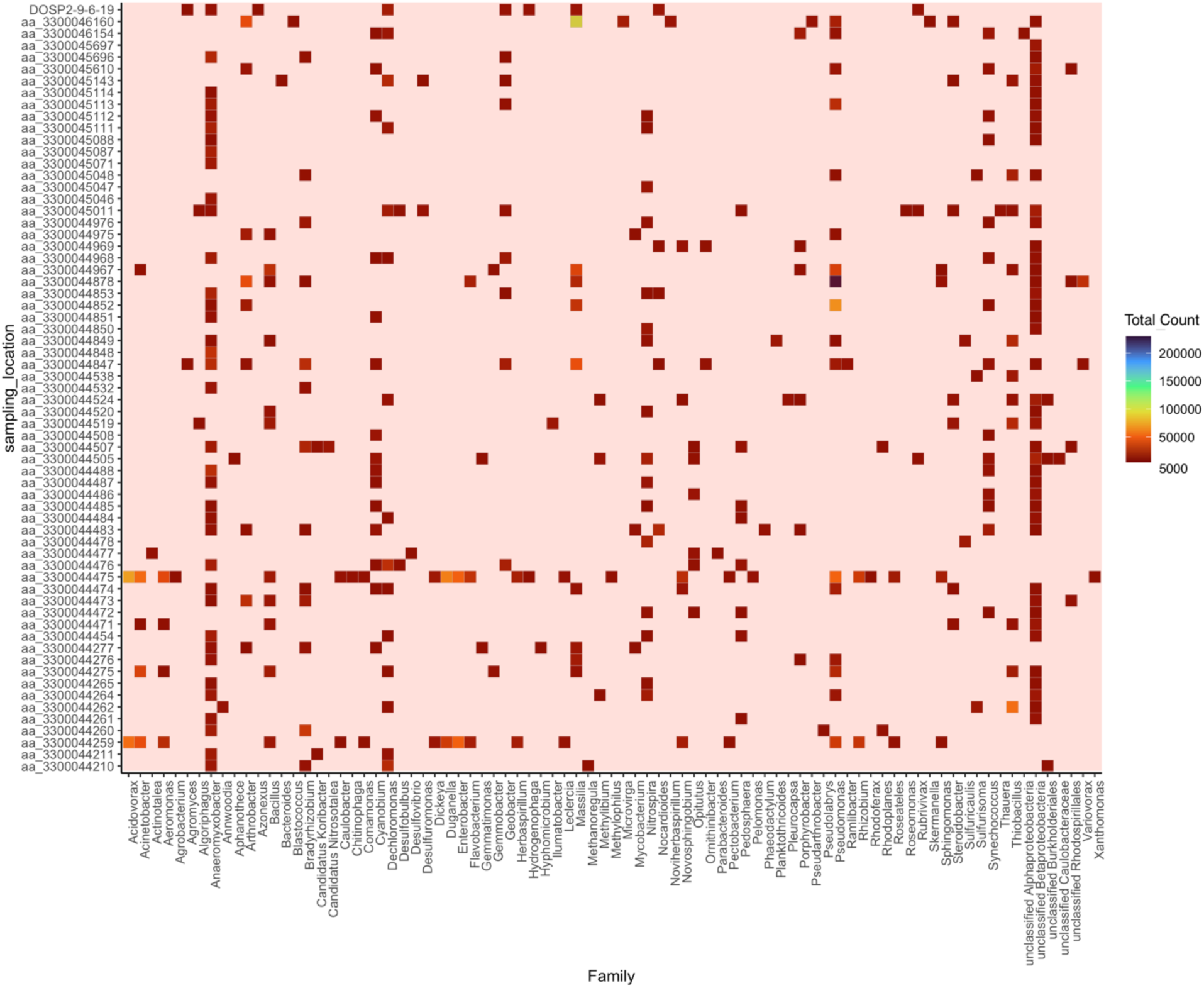
Bacterial genera per sampling site as identified by metagenomic sequences. Results were filtered for 5000 hits and greater to make a most commonly occurring heatmap.

